# Novel insights into *N*-glycan fucosylation and core xylosylation in *C. reinhardtii*

**DOI:** 10.1101/782292

**Authors:** Anne Oltmanns, Lara Hoepfner, Martin Scholz, Karen Zinzius, Stefan Schulze, Michael Hippler

## Abstract

*Chlamydomonas reinhardtii N*-glycans carry plant typical β1,2-core xylose, α1,3-fucose residues as well as plant atypical terminal β1,4-xylose and methylated mannoses. In a recent study, XylT1A was shown to act as core xylosyltransferase, whereby its action was of importance for an inhibition of excessive Man1A dependent trimming. *N*-Glycans found in a XylT1A/Man1A double mutant carried core xylose residues, suggesting the existence of a second core xylosyltransferase in *C. reinhardtii*. To further elucidate enzymes important for *N*-glycosylation, novel single knockdown mutants of candidate genes involved in the *N*-glycosylation pathway were characterized. In addition, double, triple and quadruple mutants affecting already known *N*-glycosylation pathway genes were generated. By characterizing *N*-glycan compositions of intact *N*-glycopeptides from these mutant strains by mass spectrometry, a candidate gene encoding for a second putative core xylosyltransferase (XylT1B) was identified. Additionally, the role of a putative fucosyltransferase was revealed. Mutant strains with knockdown of both xylosyltransferases and the fucosyltransferase resulted in the formation of *N*-glycans with strongly diminished core modifications. Thus, the mutant strains generated will pave the way for further investigations on how single *N*-glycan core epitopes modulate protein function in *C. reinhardtii*.

**Significance Statement:** Our data provide novel insights into the function of XylT1B and FucT in *C. reinhardtii* as *N*-glycan core modifying enzymes. In the course of our study, different mutants were created by genetic crosses showing either varying or a lack of *N*-glycan core modification, enabling comparative analyses in relation to single *N*-glycan core epitope and overall protein function in *C. reinhardtii*.

## Introduction

*N*-glycosylation is one of the major post-translational modifications of proteins in eukaryotes. It starts in the endoplasmic reticulum (ER) with the co-translational transfer of glucose_3_mannose_9_*N*-acetylglucosamine_2_ from a lipid-linked oligosaccharide precursor onto the asparagine of the consensus sequence N-X-S/T (where X may be any amino acid except proline) by the oligosaccharyl transferase complex (OST). Subsequently, two glucose residues are removed and the protein undergoes folding with help of the calnexin/calreticulin cycle. Hereupon, the last glucose and one mannose are excised and the *N*-glycoprotein is guided into the Golgi apparatus. While all *N*-glycosylation steps in the ER are highly conserved among most eukaryotes, the following maturation steps are highly dependent on organism and cell-type specific expression levels of glycosidases and glycosyltransferases. In fact, the high diversity of so-called complex type or paucimannosidic *N*-glycans in plants is a result of the manifold Golgi enzyme repertoire among different species. Typical *N*-glycan modifications absent in mammals but found in vascular plants include β1,2-xylose and α1,3-fucose. Furthermore, *N*-glycans in plants can be terminally capped by β1,3-galactose and α1,4-fucose, a structure referred to as Lewis^a^ epitope.

Although the essential role of *N*-glycosylation on protein structure and function is widely accepted, only little is known about the physiological consequences of altered *N*-glycan structures in plants (Nagashima et al., 2018). Studies in *Arabidopsis thaliana* (*A. thaliana*) suggest that *N*-glycosylation is mostly essential for protein targeting and proper protein folding in the ER, since underglycosylation leads to an accumulation of unfolded proteins in the ER lumen (Nagashima et al., 2018). Furthermore, the depletion of multiple OST subunits in *A. thaliana* often results in lethality (Koiwa et al., 2003). In contrast, in most cases plant viability does not seem to depend on *N*-glycan maturation steps in the Golgi (Strasser, 2016). Instead, mutations shifting the *N*-glycan complexity towards the oligmannosidic type, such as *xylT fucTab, cgl1* (Kang et al., 2008), *mns1 mns2* (Liebminger et al., 2009, Liu et al., 2018) and *hgl1 fucTab* (Kaulfurst-Soboll et al., 2011) were found to enhance salt sensitivity and to impact root growth in *A. thaliana*. Also, in *Oryza sativa* (*O. sativa*), knockout of the single copy of a xylosyltransferase (XylT) only affected vegetative growth at low-temperature (Takano et al., 2015).

Among the few microalgae in which *N*-glycosylation has been studied, the process is best characterized in *Chlamydomonas reinhardtii*. Recent studies revealed a unique linear structure, in which *N*-glycans are synthesized in a GnTI-independent manner but still harbor β1,2-xylose and fucose at the core (Mathieu-Rivet et al., 2013, Vanier et al., 2017). Interestingly, and uncommon for plants, *N*-glycans were shown to carry a second, terminal xylose and modifications of mannose residues with one 6-*O*-methylation. Furthermore, two *N*-glycan processing enzymes, Man1A and XylT1A, identified in an *in silico* analysis, were characterized in a recent study involving corresponding insertional mutants (Figure 1a; Schulze et al., 2018). Hereby, XylT1A was identified as β1,2-core XylT. In addition, knockout of XylT1A led to an increased trimming of *N*-glycans. Since this phenotype was abolished by the depletion of both enzymes in the same strain, an involvement of Man1A in the trimming process was concluded. Unexpectedly, knockout of Man1A also led to a loss of 6-*O*-methylation in single- and double mutants. Strikingly, core xyloses were found in the double mutant, although XylT1A was absent. This pointed towards the existence of a second core XylT (XylT1B) which was proposed to be encoded by the gene Cre16.g678997.

**Figure 1.**
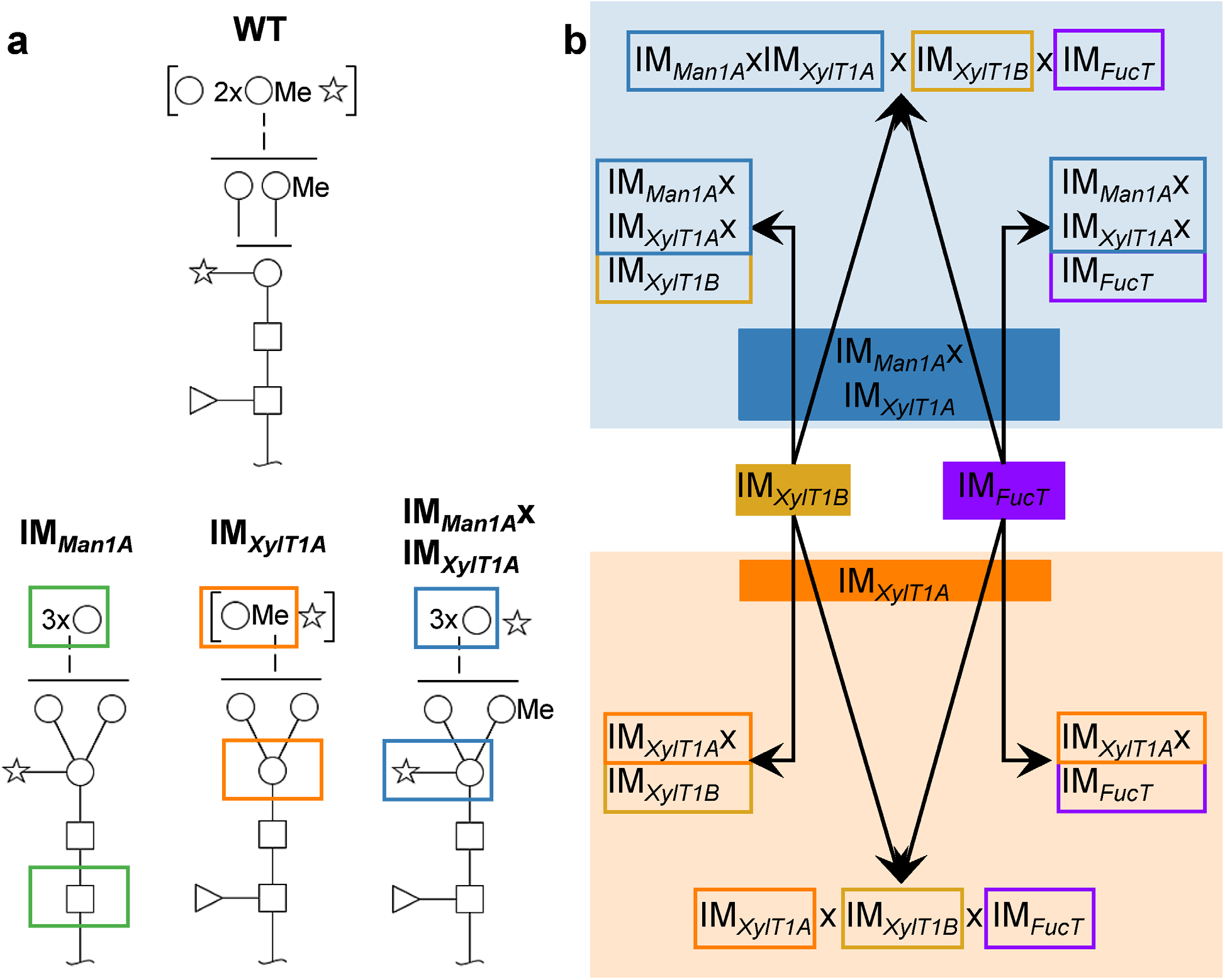
*N*-glycan compositions previously found in IM strains, WT and workflow of mutant generation within this study. *N*-glycans found in WT typically carry two xylose, one fucose and methylated hexoses. Upon knockout of XylT1A, a lack of core xylose is observed and excessive trimming occurs (orange boxes). In contrast, knockout of Man1A is causing less methylation and a partial lack of terminal xylose (green boxes). However, depletion of both enzymes in the same strain led to *N*-glycans of normal length carrying a core xylose (blue boxes) (Schulze et al. 2018). Monosaccharides depicted above the solid horizontal line can be bound to any subjacent residue, to monosaccharides connected by a dashed line or to monosaccharides within the same bracket. Monosaccharide symbols follow the Symbol Nomenclature for Glycans (a) (Varki et al. 2009). IM candidates for putative FucT and XylT1B were purchased at the CLiP library (Li et al. 2016) and successively crossed into XylT1A or Man1A/XylT1A depleted backgrounds. Additionally, mutants lacking all three putative *N*-glycan core modifying enzymes were generated (b).

To this end no physiological studies on the effect of altered *N*-glycan structures in microalgae have been reported. Since many *N*-glycosylated proteins in *C. reinhardtii* are secreted and involved in nutrient acquisition (e.g. Fe-assimilatory proteins 1 and 2, Carbonic anhydrase 1), membrane stemming ion channels (e.g. polycystic kidney disease 2; PKD2) or proteins involved in cell gliding (e.g. FMG1-B and FAP113), it is of great interest to elucidate the role of single *N*-glycan epitopes on cell physiology and protein function (Mathieu-Rivet et al., 2013, Bloodgood, 2009, Kamiya et al., 2018). This drives the need for mutants with specific lack of *N*-glycan moieties, such as e.g. core xylose or fucose.

In this study, we aimed to elucidate the identity of all enzymes involved in *N*-glycan core modification in *C. reinhardtii*. In addition to a XylT1B deficient mutant, a putative fucosyltransferase (FucT) mutant, Cre18.g749697, was analyzed. To verify the role of these enzymes, insertional mutants (IM) of the two putative glycosyltransferases were obtained from the Chlamydomonas Library Project (CLiP) (Li et al., 2016) and double, triple and quadruple mutants were generated by genetic crossing of the mutant strains. The comparative analyses of *N*-glycan compositions revealed substrate specificity of XylT1B and FucT.

## Results

To analyze the role of the putative XylT1B and FucT, insertional mutants were obtained from the CLiP (Li et al., 2016). These are characterized by the insertion of DNA cassettes encoding for a paromomycin resistance as well as for two terminator sequences (directed in both orientations) into the genomic regions of interest, hereby interrupting mRNA transcription. In addition to the single mutants, double-, triple- and quadruple mutants were generated by genetic crossings (Figure 1b). Strains derived from single cell colonies were checked by PCR for insertional cassettes (Figure S1) and Parallel Reaction Monitoring (PRM) measurements were employed to quantify relative protein abundances of Man1A, XylT1A and XylT1B (Table 1; Figure S2a). For all mutants carrying insertional cassettes in respective genes, protein amounts were below the detection limits. Since no FucT peptides were reproducibly detected in WT, reverse transcription polymerase chain reaction (RT-PCR) analysis was applied to confirm the gene knockdown on transcriptional level. Notably the insertional cassette is located close to the 3’ UTR in all strains stemming from IM*_FucT_*, yet interrupting the predicted catalytic Glycosyltransferase family 10 domain. Therefore, upstream and downstream mRNA regions were analyzed via RT-PCR. All strains carrying an insert in *fucT* were showing slightly diminished mRNA amounts prior to the insertion site, while mRNA following insertion site was drastically reduced (Figure S2b).

**Table 1.** Assessment of enzyme levels in IM strains confirms successful knockdown of corresponding genes. Protein levels of Man1A, XylT1A and XylT1B were quantified by PRM measurements. Since no peptide was reliably detected for FucT, mRNA analyses were carried out. Empty cells indicate presence of the enzyme at WT level. Data summarized here can be found in Figure S2 and Supplemental Data 1.

| IM strain | PRM (protein quantification) |  |  | mRNA analysis |
| --- | --- | --- | --- | --- |
|  | knockdown of *man1A* | knockdown of *xylT1A* | knockdown of *xylT1B* | knockdown of *fucT* |
| IM<sub>FucT</sub> |  |  |  | X |
| IM<sub>XylT1B</sub> |  |  | X |  |
| IM<sub>XylT1A</sub>XIM<sub>FucT</sub> |  | X |  | X |
| IM<sub>XylT1AB</sub> |  | X | X |  |
| IM<sub>Man1A</sub>XIM<sub>XylT1A</sub>XIM<sub>FucT</sub> | X | X |  |  |
| IM<sub>Man1A</sub>XIM<sub>XylT1AB</sub> | X | X | X | X |
| IM<sub>XylT1AB</sub>XIM<sub>FucT</sub> |  | X | X | X |
| IM<sub>Man1A</sub>XIM<sub>XylT1AB</sub>XIM<sub>FucT</sub> | X | X | X | X |

After verification of the mutants, tryptic *N*-glycopeptides of proteins secreted into the medium were analyzed by mass spectrometry. The measurements employed in-source collision induced dissociation (IS-CID), in which ion acceleration within the ion source region of the mass spectrometer leads to an incomplete fragmentation of glycan chains (Hsiao and Urlaub, 2010). The resulting glycopeptide ions with varying glycan length are then analyzed by a conventional survey scan (MS1). Using the “mass tags” option, ions that differ in mass by 203 or 406 Da, corresponding to one or two *N*-acetylhexosamine (HexNAc) residues, respectively, are further selected for higher-energy collisional dissociation (HCD) fragmentation (MS2). After data acquisition, peptide identification by common database search engines is carried out with HexNac and HexNAc(2) set as variable modifications. In addition, with help of the in-house developed python-based tool SugarPy (Schulze et al., 2017, Schulze et al., 2018), the *N*-glycopeptide fragment ion series observed in MS1 can be analyzed to reconstruct the *N*-glycan composition (examples are given in Figure S3). By gaining information about both, peptide and attached carbohydrate moiety, *N*-glycan compositions of specific *N*-glycosites found in two strains can be directly compared, i.e. parameters such as differences in *N*-glycan length and differential number of pentose (Pent) or deoxyhexose (dHex) attached to the same *N*-glycopeptide in two strains can be directly assessed (for examples of these calculations see the Material and Methods section). Thereby, providing insights into the function of the analyzed glycosyltransferases.

### Role of FucT

Tryptic *N*-glycopeptides stemming from supernatant (SN) samples of the single mutant IM_*FucT*_ were analyzed by mass spectrometry (Figure S4a). Strikingly, the knockdown of FucT did not yield a lack of dHex when comparing *N*-glycosites to WT (Figure S4e). Instead, *N*-glycans were slightly reduced in length, defined as the sum of hexose (Hex) and methylated hexose (MeHex). Consequently, a knockdown of the putative fucosyltransferase alone does not have an impact on the fucose transfer onto *N*-glycans, but apparently on *N*-glycan length.

In order to analyze the role of FucT in a different background, the triple mutant IM*_Man1A_*xIM*_XylT1A_*xIM*_FucT_* was created by genetic crossing. In the following, *N*-glycan compositions of the triple mutant IM*_Man1A_*xIM*_XylT1A_*xIM*_FucT_* were compared to *N*-glycans found in IM*_Man1A_*xIM*_XylT1A_* (Figure 2a and Figure S5). *N*-Glycans attached to peptides found in both strains showed a similar length and number of Pent. Meanwhile, a strong decrease in the number of *N*-glycosites carrying dHex was apparent for IM*_Man1Ax_*IM*_XylT1A_*xIM*_FucT_* (Figure 2b).

In summary, *fucT* expression affects Man1A-dependent trimming. Furthermore, fucose transfer is drastically reduced after *fucT* knockdown in Man1A-/XylT1A-depleted genetic backgrounds.

**Figure 2.**
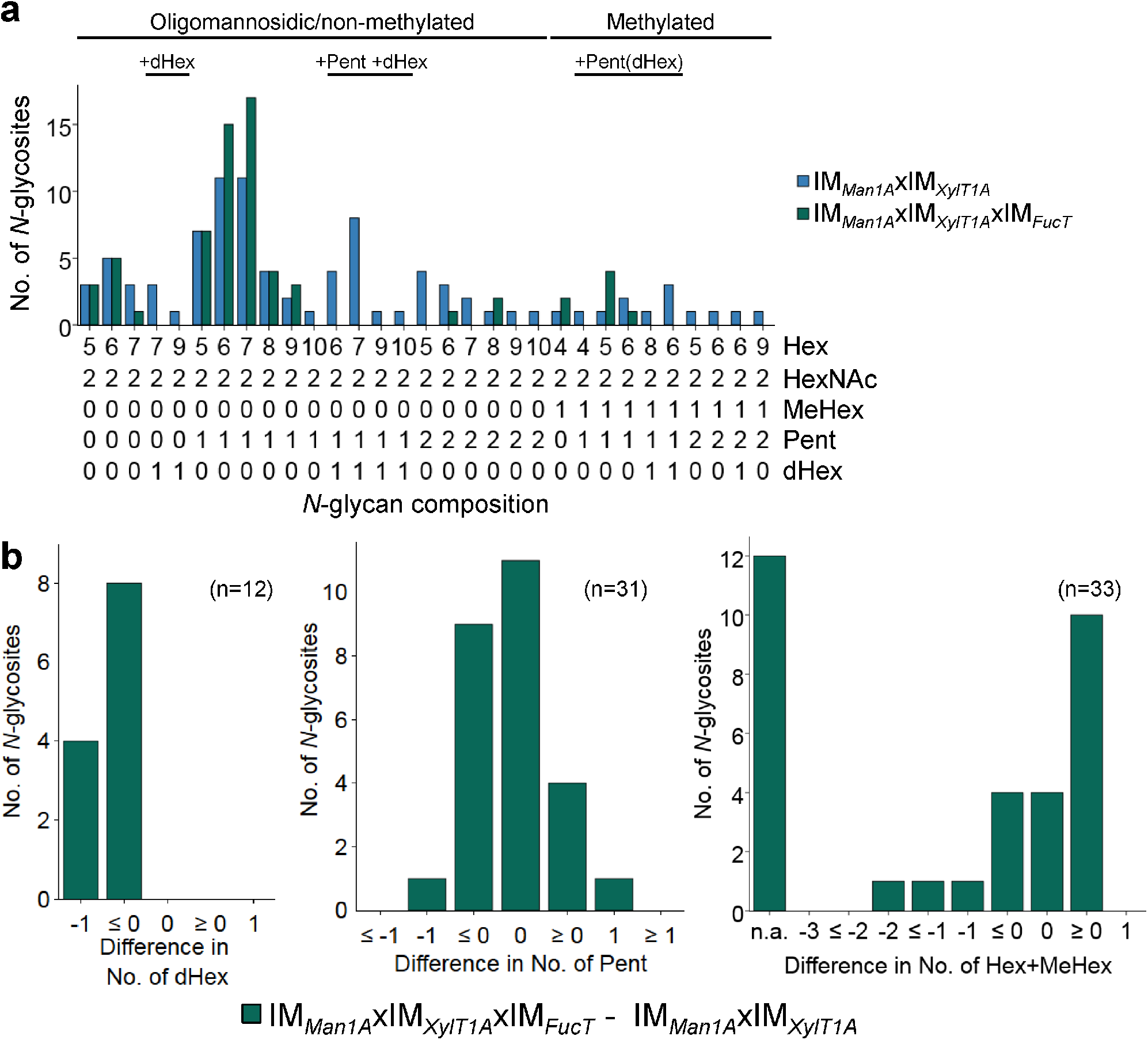
*N*-glycan compositions of IM*_Man1A_*xIM*_XylT1A_* and IM*_Man1A_*xIM*_XylT1A_*xIM*_FucT_* differ in dHex levels. a, For all identified *N*-glycan compositions, the number of *N*-glycosites harboring this glycan is shown for IM*_Man1A_*xIM*_XylT1A_* (blue) and IM*_Man1A_*xIM*_XylT1A_*xIM*_FucT_* (green). The *N*-glycan complexity is increasing from left (oligomannosidic, not methylated) to right (decorated, methylated). *N*-glycan compositions were grouped according to the presence of Pent and/or dHex (optional for sugars written in parenthesis). Only *N*-glycosites identified in both strains were taken into account (n=33). Peptide sequences and *N*-glycan compositions attached are listed in Supplemental Data 2. b, Differences in the number of dHex (left), Pent (middle) and Hex+MeHex (right) for *N*-glycosites found in both strains. *N*-glycosites carrying no dHex (left) or Pent (middle) in both strains were excluded. The legends indicate the total number of *N*-glycosites compared. Some *N*-glycosites harboring multiple *N*-glycoforms could not be assigned to one of the categories (n.a.); for dHex n.a.=0 and for Pent n.a.=5.

### XylT1B cannot act on excessively trimmed *N*-glycans

The existence of a second core XylT (XylT1B) had been proposed earlier when core xylose was found attached to *N*-glycans in the double mutant IM*_Man1A_*xIM*_XylT1A_*. At the same time, this second XylT seemed unable to act in IM*_XylT1A_*, as a clear reduction in core Pent was seen in the single IM strain. Therefore, it had been hypothesized that excessive trimming would prevent core modification by XylT1B (Schulze et al., 2018). The comparison between the putative XylT1B single mutant and WT, revealed no clear differences in terms of *N*-glycan composition (Figure S4b). The overall distribution of *N*-glycans resembled the composition of WT: methylated *N*-glycans as well as *N*-glycans carrying up to two Pent and one dHex were identified (Figure S4e). At the same time, the amount of Pent or dHex attached to *N*-glycans on the same *N*-glycosites was not altered in comparison to the WT (Figure S4f). These findings suggest that either XylT1B is not involved in *N*-glycosylation or that a lack of XylT1B is compensated by XylT1A. To test the latter hypothesis, several IM strains were created by genetic crossing: IM*_Man1A_*xIM*_XylT1AB_* (notably, both XylTs are affected here), IM*_XylT1AB_* (a double mutant of the single XylT IM strains) and IM*_XylT1A_*xIM*_FucT_*.

When comparing IM*_Man1A_*xIM_*xylT1A*_ to IM*_Man1A_*xIM*_xylT1AB_*, mass spectrometric analyses revealed a drastic decrease in the number of Pent per *N*-glycopeptide while length and degree of dHex are not altered (Figure 3, Figure S6d). Although no *N*-glycan linkages can be determined via the mass spectrometry measurements conducted, the presence or absence of certain masses can be compared between two strains. Peaks corresponding to “peptide+HexNAc(2)+Hex+Pent”, indicated the existence of a core xylose. While found in WT spectra, these peaks were absent in IM*_Man1A_*xIM*_XylT1AB_* (Table S1), thus suggesting that mainly core instead of terminal xylose were lost in the mutant.

**Figure 3.**
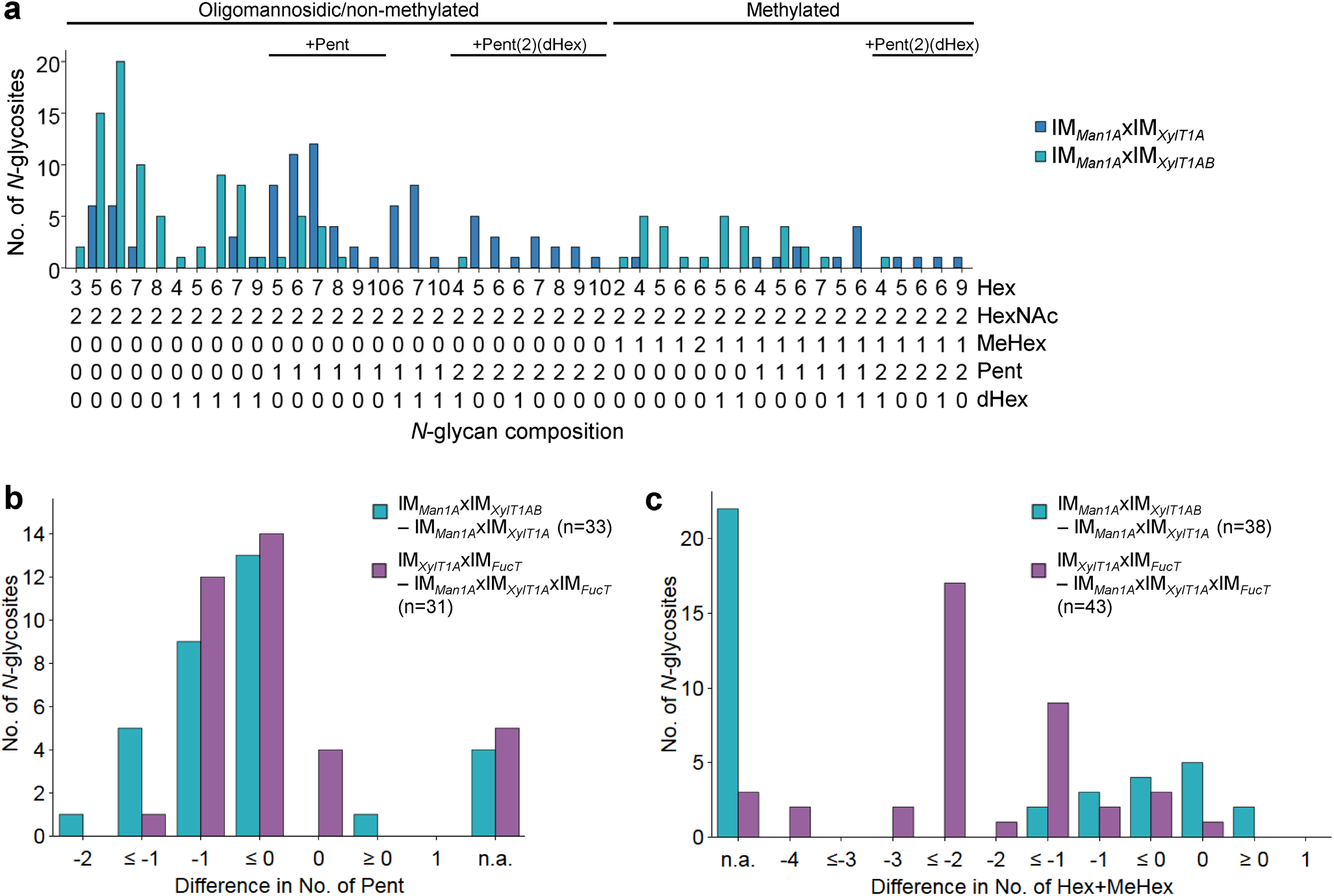
Knockdown of XylT1B in IM*_Man1A_*xIM*_XylT1A_* leads to lack of Pent. a, For all identified *N*-glycan compositions, the number of *N*-glycosites harboring this glycan is shown for IM_*Man1A*_xIM*_XylT1A_* (dark blue) and the triple mutant IM*_Man1A_*xIM*_XylT1AB_* (light blue). The *N*-glycan complexity is increasing from left (oligomannosidic, not methylated) to right (decorated, methylated). *N*-glycan compositions were grouped according to the presence of Pent and/or dHex (optional for sugars written in parenthesis). Only *N*-glycosites identified in both strains were taken into account (n=38). Peptide sequences and *N*-glycan compositions attached are listed in Supplemental Data 2. Differences in the number of Pent (b) and in *N*-glycan length (c) for *N*-glycosites found in both strains were calculated as depicted in the legend. *N*-glycosites carrying no Pent (b) both strains were excluded. The legends indicate the total number of *N*-glycosites compared. Some *N*-glycosites harboring multiple *N*-glycoforms could not be assigned to one of the categories (n.a.).

When assuming that the putative XylT1B can act as a core XylT, *N*-glycans found in IM*_XylT1A_* and IM*_XylT1AB_* should not differ as IM*_XylT1A_* is already depleted in core xylose (Schulze et al., 2018). Indeed, the overall *N*-glycan compositions did not differ and no reduction in the amount of Pent was observed when comparing the two mutant strains (Figure S7).

To confirm the hypothesis regarding substrate specificity of XylT1B towards WT length *N*-glycans (Schulze et al., 2018), *N*-glycans found in IM*_XylT1A_*xIM*_FucT_* and IM*_Man1A_*xIM*_XylT1A_*xIM*_FucT_* were compared. Notably, while both strains expressed XylT1B (Table 1) and were devoid of dHex to the same degree (Figure S6d), *N*-glycans found in IM*_XylT1A_*xIM*_FucT_* were excessively trimmed and therefore significantly shorter than *N*-glycans from IM*_Man1A_*xIM*_XylT1A_*xIM*_FucT_* (Figure 3c). At the same time, *N*-glycans in IM*_XylT1A_*xIM*_FucT_* carried fewer (core) Pent than *N*-glycans found in the triple mutant (Figure 3b, Table S1). Consequently, *N*-glycan compositions obtained for these two mutants strengthen the hypothesis that XylT1B cannot act, when *N*-glycans are excessively trimmed.

### Generation of strains devoid of *N*-glycan core modifications

To analyze, whether a knockdown of FucT, XylT1A and XylT1B would be sufficient to eliminate all *N*-glycan core modifications in *C. reinhardtii*, another triple mutant (IM*_XylT1AB_*xIM*_FucT_*) and a quadruple mutant (IM*_Man1A_*xIM*_XylT1AB_*xIM*_FucT_*) were generated by genetic crosses. By considering *N*-glycosites found in WT and IM*_XylT1AB_*xIM*_FucT_*, a significant shift in the overall *N*-glycan compositions towards less complex *N*-glycans was seen (Figure 4a). In line with this, less *N*-glycosites carrying core Pent or dHex (Figure 4c; Table S1) were identified in the triple mutant. Indeed, the same trend was seen when considering all *N*-glycosites found (Figure S8a). Also, when comparing the quadruple mutant with the WT, no considerable amounts of core Pent or dHex were found (Figure 4c; Table S1), while the majority of *N*-glycosites harbored oligomannosidic *N*-glycans in IM*_Man1A_*xIM*_XylT1AB_*xIM*_FucT_* in contrast to complex *N*-glycans in the WT (Figure 4b).

**Figure 4.**
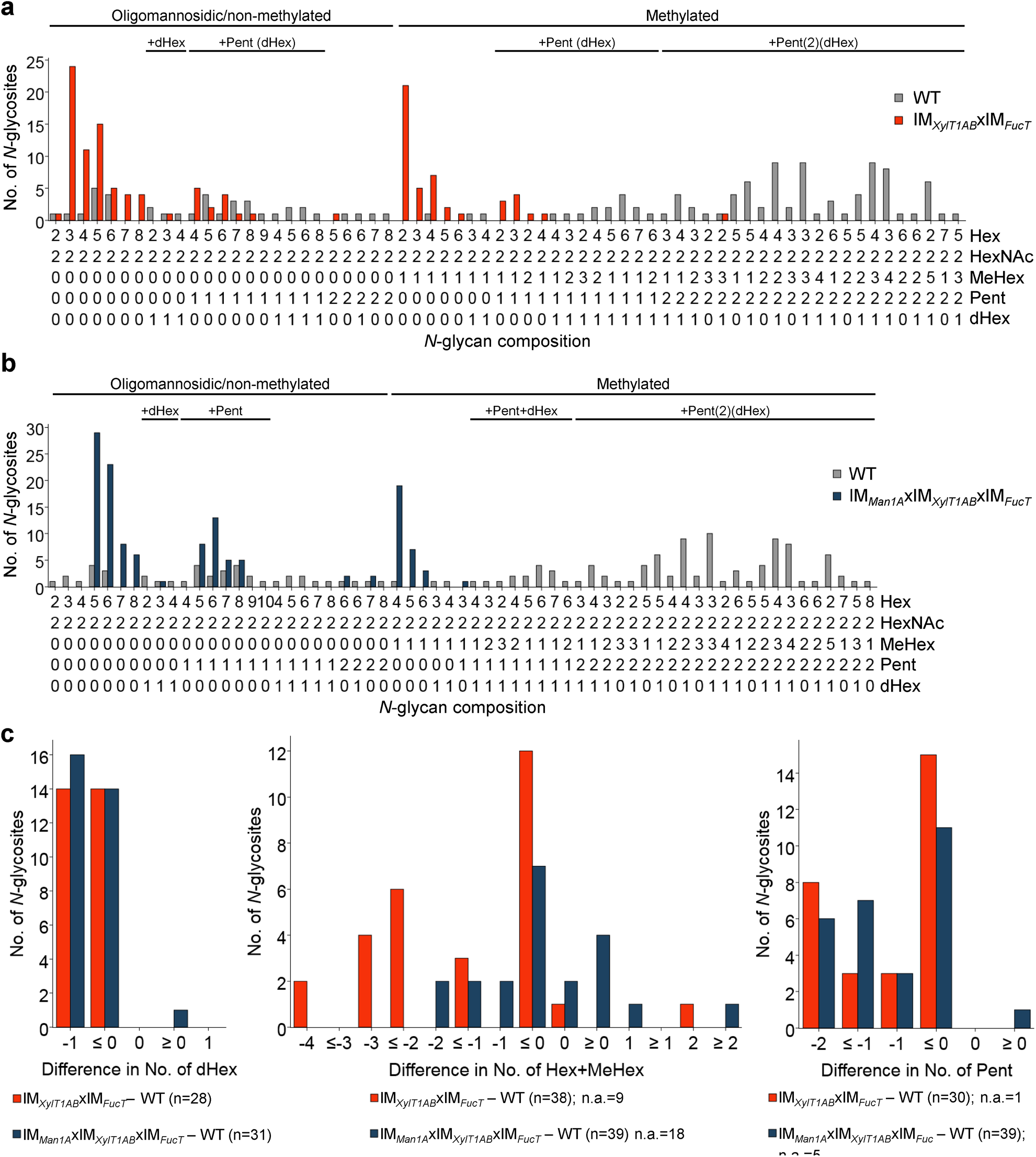
Depletion of XylT1A, XylT1B and FucT leads to strains devoid of *N*-glycan core modifications. For all identified *N*-glycan compositions, the number of *N*-glycosites harboring this glycan is shown for WT (grey) and the triple mutant IM*_XylT1AB_*xIM*_FucT_* (orange) (a) and the quadruple mutant IM*_Man1A_*xIM*_XylT1AB_*xIM*_FucT_* (dark blue) (b), respectively. The *N*-glycan complexity is increasing from left (oligomannosidic, not methylated) to right (decorated, methylated). *N*-glycan compositions were grouped according to the presence of Pent and/or dHex (optional for sugars written in parenthesis). Only *N*-glycosites identified in both strains compared were taken into account (n=38 for A and n=39 for b). Peptide sequences and *N*-glycan compositions attached are listed in Supplemental Data 2. c, Differences in the number of dHex (left), *N*-glycan length, defined as the sum of Hex+MeHex (middle), and in the number of Pent (right) for *N*-glycosites found in both strains compared were calculated as depicted in the legend. *N*-glycosites carrying no dHex (left) or Pent (right) in both strains were excluded, respectively. The legends indicate the total number of *N*-glycosites compared. Some *N*-glycosites harboring multiple *N*-glycoforms could not be assigned to one of the categories (n.a.).

### Immunoblotting proves knockout of *N*-glycosylation enzymes

For an independent verification of *N*-glycan compositions derived by mass spectrometric analyses, a polyclonal anti-horseradish peroxidase (HRP) antibody was affinity purified using *A. thaliana* mutant leaf extracts (*fucTab*, *xylT* and *xylT fucTab*). Subsequently, immunoblots of supernatant (SN) proteins of *C. reinhardtii* were probed with antibody fractions binding to β1,2-xylose and α1,3-fucose residues attached to *N*-glycan cores, respectively (Figure 5 and S9). Signals obtained from *A. thaliana* mutant leaf extracts prove the specificity of affinity purified antibody fractions. Considering SN samples of *C. reinhardtii*, the same protein amounts were loaded for the different strains. The Coomassie control stain revealed differential protein ratios within the single lanes indicating differential secretion of proteins despite identical culture conditions. Therefore, changes in protein levels are attributed to to distinct *N*-glycan patterns in the strains.

**Figure 5.**
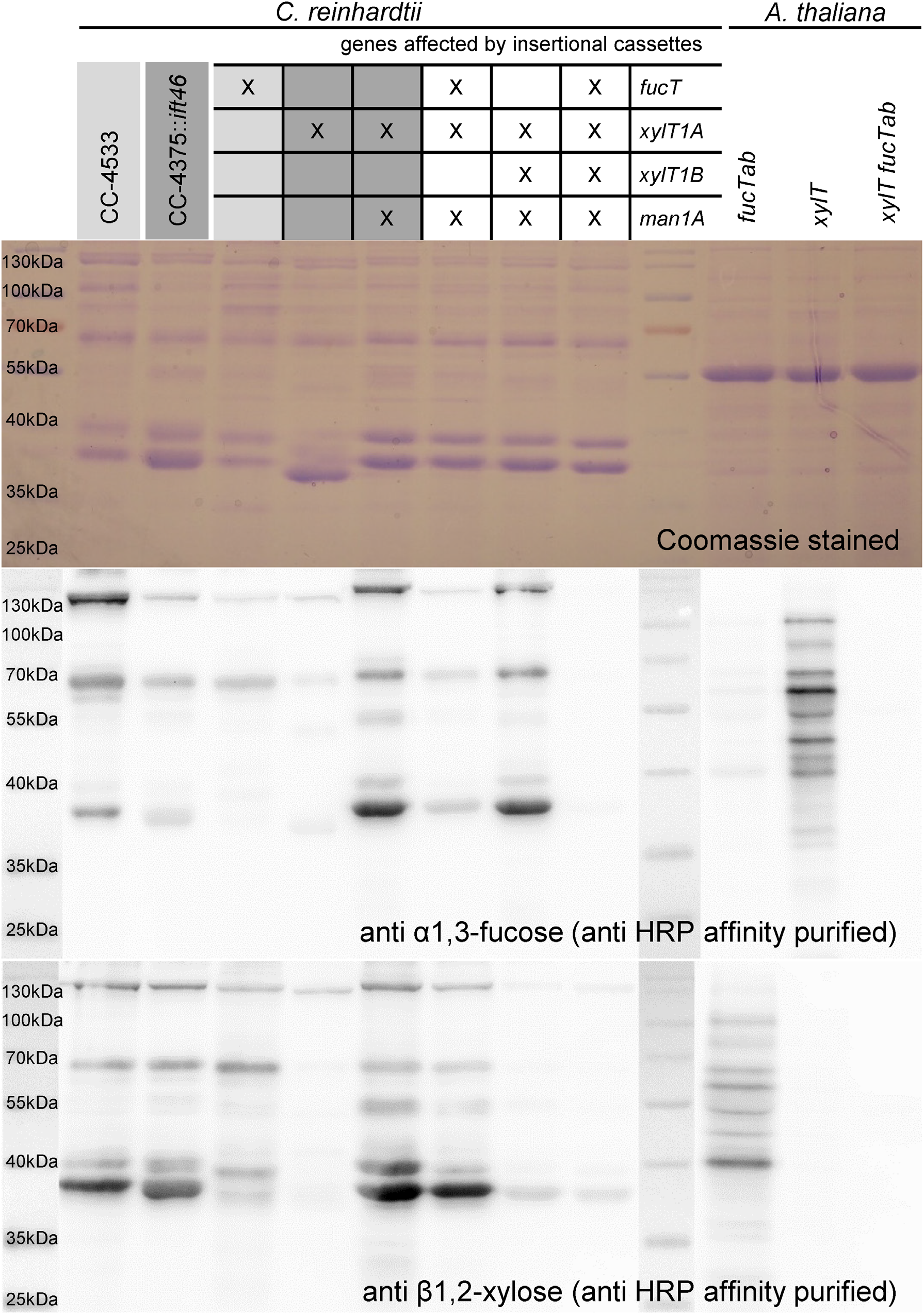
Immunoblotting confirms loss of β1,2-core xylose and α-1,3-fucose in IM strains. 20 μg of SN proteins of selected IM strains as well as of *A. thaliana* leaf extracts were separated by SDS-PAGE in triplicates and transferred to nitrocellulose membranes. Additionally, one triplicate was stained as loading control using Coomassie Brilliant Blue G (top). Membranes were incubated in affinity purified HRP antibody binding to α-1,3-fucose (middle) and β1,2-core xylose (bottom), respectively. Strains highlighted in different shades of grey indicate known corresponding WT backgrounds.

**Figure 6.**
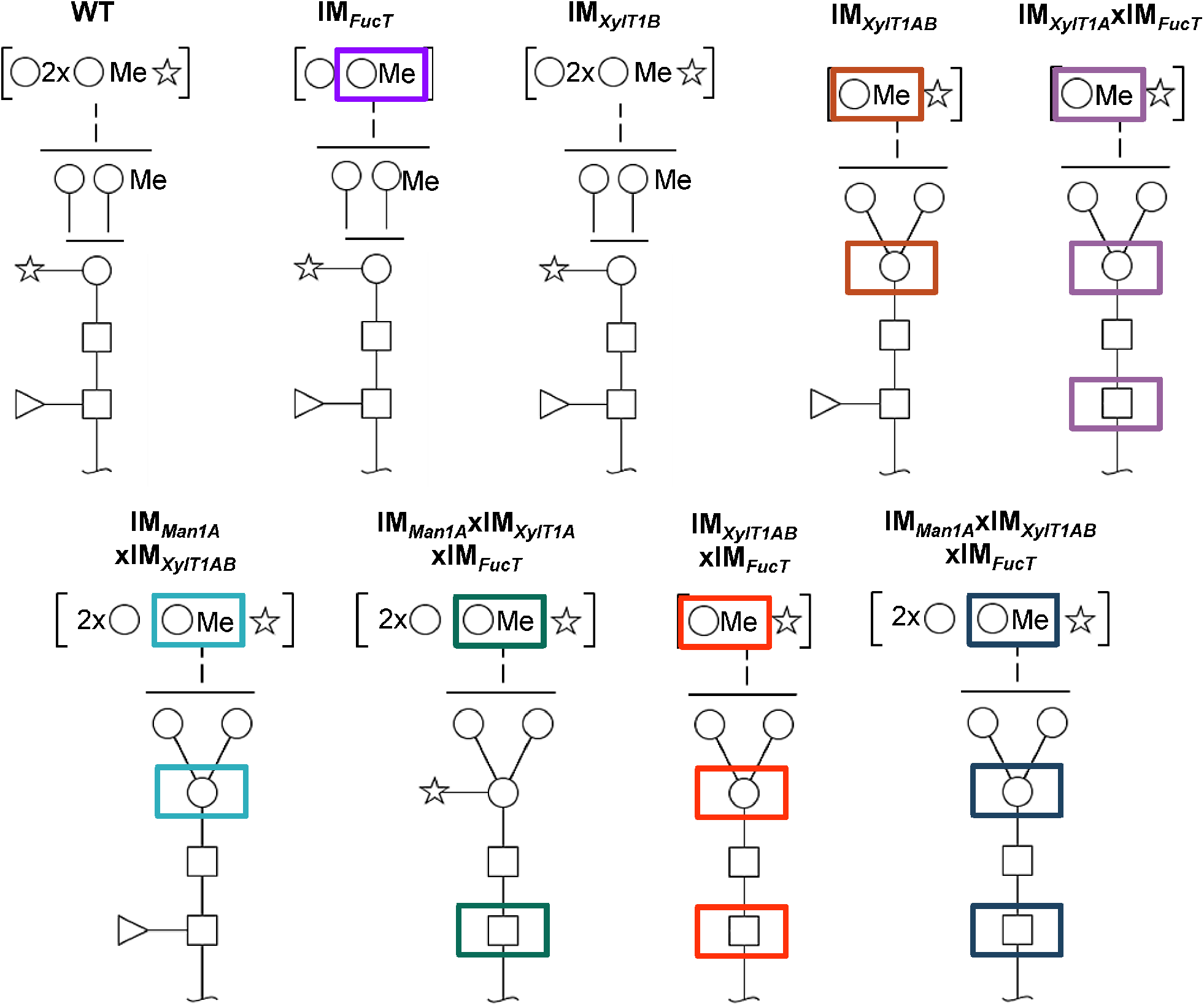
Schematic representation of characteristic N-glycan compositions found in new IM strains and WT. While knockdown of FucT leads to lack of fucose in multiple mutants when core xylose is absent, *N*-glycan length is diminished in the single IM strain. Interestingly, knockout of only XylT1B has no effect on *N*-glycan compositions indicating a minor role in the *N*-glycosylation process in comparison to XylT1A. When knocking out XylT1B in a mutant background lacking Man1A and XylT1A (leading to WT length *N*-glycans) core xyloses are not observed indicating a role of XylT1B as core XylT. As shown previously, knockout of XylT1A causes excessive Man1A dependent trimming, while knockout of Man1A leads to a lack of methylation. Monosaccharides depicted above the solid horizontal line can be bound to any subjacent residue, to monosaccharides connected by a dashed line or to monosaccharides within the same bracket. Monosaccharide symbols follow the Symbol Nomenclature for Glycans (Varki et al. 2009).

In general, both antibodies showed lower affinity towards *N*-glycoproteins secreted by WT than towards *N*-glycoproteins secreted by IM*_Man1A_*xIM*_XylT1A_*. As it was shown previously that WT *N*-glycans are highly methylated while *N*-glycans produced in IM*_Man1A_*xIM*_XylT1A_* are reduced in methylation, one possible explanation might be an altered accessibility to the *N*-glycan core epitopes (Schulze et al., 2018). While conformational changes have been shown to be of importance for antibody recognition (Kaulfurst-Soboll et al., 2011), methylation might change the overall conformation or reduce the antibody accessibility by direct shielding of the *N*-glycan core.

In regard to the anti-α1,3-fucose antibody and mutants affected in *fucT*, a minor signal decrease was observed in IM*_FucT_*, while the quadruple mutant showed only marginal residual recognition by the antibody (Figure 5). Unexpectedly, when considering mass spectrometric results which showed a clear decrease in dHex (Figure 2b), the signal strength of IM*_Man1A_*xIM*_XylT1A_*xIM*_FucT_* was comparable to CC4375*::ift46*, although decreased in comparison to IM*_Man1A_*xIM*_XylT1A_*, indicating the presence of α1,3-fucose in this mutant.

When looking at the anti-β1,2-xylose antibody recognition, low signal of IM*_XylT1A_* as well as high signal of IM*_Man1A_*xIM*_XylT1A_* and IM*_Man1A_*xIM*_XylT1A_*xIM*_FucT_* confirmed already published results (Schulze et al., 2018). When XylT1B is knocked down in IM*_Man1A_*xIM*_XylT1A_*, the antibody signal was reduced to a minimal background level indicating the loss of a core xylose, thereby confirming the loss of a core Pent as seen in mass spectrometric measurements (Figure 3b). Also, for the quadruple mutant, only minimal background signal was obtained.

## Discussion

In this work we provide new insights into the function of XylT1B and FucT in *C. reinhardtii*.

Notably, the composition of *N*-glycans observed in this study is in line with what has been described previously for *C. reinhardtii*: *N*-glycans carrying dHex, up to two Pent and methylated Hex were identified(Mathieu-Rivet et al., 2013, Schulze et al., 2018). Taking advantage of anti-β1,2-xylose and anti-α1,3-fucose specific antibodies, the immunoblot analyses confirmed the dHex attached to *N*-glycans as α1,3-fucose and one of the Pent as β1,2-core xylose. The position of the remaining Pent could not be resolved due to a lack of specific antibodies. However, newest data suggest that the additional Pent is a xylose attached to the first or second Hex of the α1,3-branch (Lucas et al. 2019).

### FucT transfers α1,3-fucose onto *N*-glycans in a core xylose dependent manner

The data presented herein indicate that FucT adds α1,3-fucose to the *N*-glycan chain. The conclusion that FucT indeed has FucT activity is revealed by mass spectrometric data as the knockdown of *fucT* in different genetic backgrounds (IM*_XylT1A_*xIM*_FucT_*, IM*_Man1A_*xIM*_XylT1A_*xIM*_FucT_*, IM*_XylT1AB_*xIM*_FucT_* and IM*_Man1A_*xIM*_XylT1AB_*xIM*_FucT_*), caused a loss of dHex. In line with mass spectrometric analysis, a strong diminishment of α1,3-fucose (corresponding to a loss of dHex) was verified by immunoblotting for IM*_XylT1A_*xIM*_FucT_*, IM*_XylT1AB_*xIM*_FucT_* and IM*_Man1A_*xIM*_XylT1AB_*xIM*_FucT_*(Figure 5, Figure S9). The comparison of mass spectrometric data and immuno-blotting results indicated slight differences in the presence and absence of α1,3-fucose. In this line, mass spectrometric analyses for IM*_Man1A_*xIM*_XylT1A_*xIM*_FucT_* indicated the almost complete absence of dHex (Table S1) whereas immunoblot results implied the presence of α1,3-fucose (Figure 5). Similarly, mass spectrometric analyses predicted no changes in fucosylation in IM*_FucT_* (Figure S4f), while immunoblotting showed a slightly decreased signal. These differences could be explained by the fact that immunoblotting detects proteins from the whole secretome, while mass spectrometric measurements describes the *N*-glycan composition of individual peptides. Moreover, it has to be stated that mass spectrometric data presented in this study are not quantitative. To account for those biases, both methods were employed and, overall, mass spectrometric and immunoblot analyses are in accordance.

The second interesting finding was that a complete lack of α1,3-fucose occurred only in mutants additionally depleted in core xylose (IM*_XylT1A_*xIM*_FucT_*, IM*_XylT1AB_*xIM*_FucT_* and IM*_Man1A_*xIM*_XylT1AB_*xIM*_FucT_*). Consequently, data obtained for multiple mutants lacking FucT suggest an interconnection of fucosylation and core-xylosylation: when knocking down FucT in core xylose deficient backgrounds, core fucose is equally lost. This might be caused either by a strictly sequential enzymatic action of XylT and FucT or a physical interaction between several enzymes involved in *N*-glycosylation. Neither of the two possibilities can be ruled out by data obtained in this study. A third possible explanation for the presence of fucose residues in IM*_FucT_* as well as in IM*_Man1A_*xIM*_XylT1A_*xIM*_FucT_* would be the presence of a second FucT in *C. reinhardtii* compensating the lack of enzyme in both mutants. According to immunoblot results, this enzyme would require *N*-glycan substrates carrying core xylose residues and therefore would not be active in core xylose lacking IM strains analyzed in this study. Although there is no second α1,3-FucT predicted in the genome, several O-FucTs (Cre08.g361600, Cre03.g206705, Cre09.g388134 and Cre13.g588850) with high homology scores for different *A. thaliana* O-FucTs as well as few α1,6-FucTs (Cre08.g364351, Cre10.g449050 and Cre18.g749047) are predicted in the *C. reinhardtii* genome.

### XylT1B transfers β1,2-xylose onto *N*-glycans

As shown for FucT, five insertional mutant strains of XylT1B were analyzed, which allowed insights into the role of XylT1B in the *N*-glycosylation process (IM*_XylT1B_*, IM*_XylT1AB_*, IM*_Man1A_*xIM*_XylT1AB_*, IM*_XylT1AB_*xIM*_FucT_* and IM*_Man1A_*xIM*_XylT1AB_*xIM*_FucT_*). Initially, the presence of a second core XylT had been proposed, when *N*-glycans in a Man1A/XylT1A doublemutant were found to carry core xylose residues. When XylT1B was knocked out in addition to XylT1A and Man1A, *N*-glycans lost one Pent as revealed by mass spectrometric analyses. Immunoblotting results confirmed that β1,2-xylose was lost by knocking down XylT1B in the double mutant instead of a β1,4-xylose, which is attached to an outer mannose residue in *C. reinhardtii* (Mathieu-Rivet et al., 2013). Therefore, it can be concluded that XylT1B is a core XylT. However, it cannot be excluded that XylT1B also possesses additional XylT activity thereby possessing a higher flexibility in its acceptor substrate specificity.

Additionally, data presented here confirm the previous hypothesis, that XylT1B cannot act on excessively trimmed *N*-glycans (Schulze et al., 2018), while it was furthermore shown, that its lack can be compensated by XylT1A as seen in IM*_xylT1B_*.

### IM*_XylT1AB_*xIM*_FucT_* and IM*_Man1A_*xIM*_XylT1AB_*xIM*_FucT_* are depleted in β1,2-xylose and α1,3-fucose

As proof of principle, triple and quadruple mutants were created to analyze, whether knockdown of all FucT and XylT analyzed so far would result in *N*-glycans devoid of core xylose and fucose residues. In fact, those mutants (IM*_XylT1AB_*xIM*_FucT_* and IM*_Man1A_*xIM*_XylT1AB_*xIM*_FucT_*) did not carry significant amounts of core xylose or fucose residues as shown by immunoblot and mass spectrometric results and mainly differed in their *N*-glycan length (Figure 4 and 5). Residual core-fucose and -xylose residues found in those two mutants can probably be attributed to residual enzymatic activity, since the study was carried out using knockdown instead of knockout mutants. Although the existence of further *N*-glycan core modifying enzymes cannot be fully excluded, these two mutants already represent a great potential for assessing the impact of differential *N*-glycan core modification on cell physiology.

In conclusion, our data provide novel insights into the function of XylT1B and FucT in *C. reinhardtii*. Additionally, enzyme substrate specificities and implications on the overall *N*-glycosylation process were described. In the course of our study, a set of different mutants showing varying *N*-glycan core modifications has been created. This enables comparative analyses for gaining more knowledge on the function of single *N*-glycan core modifications when it comes to overall protein function in *C. reinhardtii*.

## Experimental procedures

### Growth conditions

All *C. reinhardtii* strains were grown photoheterotrophically in TAP medium at 25 °C and 20 μE m^-2^ s^-1^ (low light; LL) or 80μE m^-2^ s^-1^ (normal light; NL), either as liquid cultures shaking at 120 r.p.m. or on TAP plates containing 1.5 % agar. For quantitative proteomics experiments, isotopic ^14^N and ^15^N labeling was performed using ^14^N and ^15^N ammonium chloride, respectively.

### IM strains used

Insertional mutants with the accession numbers LMJ.RY0402.049160 (Cre18.g749697) and LMJ.RY0402.118417 (Cre16.g678997) were purchased at the CLiP (Li et al., 2016) library.

After streaking the cells into single colonies, strains were tested by PCR for insertion of the AphVIII fragment as recommended. Primers used can be found in Table S2. Furthermore three strains (IM*_Man1A_*, IM*_XylT1A_* and IM*_Man1A_*xIM*_XylT1A_*), described in (Schulze et al., 2018) were used for genetical crosses and for comparative reasons on DNA level. The WT strain, used for protein quantification by Parallel-Reaction-Monitoring, Western Blotting and PCR analysis, can be considered as a genetical mixture between CC-4375 and CC-4533. *N*-glycan data for WT is partly taken from Supplemental Material published in (Schulze et al., 2018) and partly from a WT strain, CC-4533 (which represents the background strain of the IM from the CLiP library), analyzed in this study. For comparison of new IM strains with data from IM*_Man1A_*, IM*_XylT1A_* or IM*_Man1A_*xIM*_XylT1A_* Supplemental Material from (Schulze et al., 2018) was used.

### Quantification of Man1A and XylT1A expression levels by Parallel-Reaction-Monitoring

Cells were grown under LL conditions in ^14^N and ^15^N TAP media. The experiment was performed as label swap experiment. Cell amounts corresponding to 5 μg chlorophyll were mixed (IM strain and WT) and pelleted by centrifugation (5,000 g, 5 minutes). Cells were lysed in 100mM Tris-HCl buffer pH=7.6 containing 2% SDS, 1 mM PMSF and 1 mM Benzamidine using a sonication bath for 5 minutes. After removal of cell debris, soluble proteins were subjected to a FASP protocol as described below using 0.35 μg trypsin.

The Q Exactive Plus was operated with the following PRM settings: resolution: 70,000 at *m/z* 200, AGC target: 5e4, maximum injection time: 240 ms, isolation window: 1.6 *m/z*. The gradient used for peptide elution with a flow rate of 300 nL min^-1^ is specified in Table S3. Inclusion lists compiled with Skyline (MacLean et al., 2010; version 4.1) were used for scheduled fragmentation of target peptides.

The total peak areas of a minimum of three fragment ions per peptide were determined in Skyline with manual adjustment of peak borders. In order to correct for differences in total protein levels in the samples, peak areas were multiplied by a replicate-specific normalization factor derived from the mean abundance of three mitochondrial ATPase subunit peptides. Peptide levels and standard deviations were calculated based on the peak areas of ^14^N and ^15^N labeled samples of the respective strain within the label swap. The .raw files can be accessed together with the quantification results (see Data availability).

### mRNA extraction and cDNA analysis

Cells were grown under normal light conditions. Aliquots corresponding to a chlorophyll amount of 40 μg were pelleted by centrifugation and resuspended in TRI Reagent™ Solution (Thermo Fisher Scientific). mRNA was extracted according to the manufacturer’s protocol and cDNA synthesis was performed, using the iScript™ cDNA Synthesis Kit (Bio-Rad) according to the manufacturer’s protocol. cDNA analysis was performed in standard PCR reactions, adjusting the cycle number of target proteins to tubulin expression levels.

### Protein isolation from the culture supernatant

The culture supernatant was obtained by two consecutive centrifugation steps. First, cells were pelleted at 5,000 g for 5min. Then, the SN was centrifuged at 48,000 g for 2 h at 4 °C. The resulting SN was concentrated by a factor of about 100 by centrifugation in filter units at 4 °C (Amicon ultra centrifugal filters, 15 mL, 10 kDa MWCO, Millipore). Protein concentration was determined by bicinchoninic acid assay (BCA Protein Assay Kit by Thermo Scientific Pierce). Samples were frozen in liquid nitrogen and stored at −80 °C until use.

**Protein isolation from *A. thaliana* leaf tissue** 1 mg of A. thaliana leaf tissue of *xylt, fucTab* and *xylt fucTab* (plants kindly provided by Prof. Dr. Antje von Schaewen) were ground in 450 μL of lysis buffer (50 mM Hepes, 250mM NaCl, 1 mM DTT, 1 mM PMSF, 1 mM Benzamidine, pH=7,5), heated to 65° C for 20 min and cell debris was pelleted at 14,000 g for 10 min at 4° C, respectively. The protein concentration of the clear supernatant was determined using the bicinchoninic acid assay (BCA Protein Assay Kit by Thermo Scientific Pierce).

### Affinity purification of HRP antibody

The antibody fractions of the polyclonal HRP antibody mixture (P7899, Sigma-Aldrich) binding to α1,3-fucose and anti β1,2-xylose, respectively, were isolated following Kaulfurst-Soboll et al., 2011 and Frank et al 2008. Deviating from the published protocol, approx. 500 μg of leaf extract protein were separated per SDS-PAGE and an initial HRP antibody dilution of 1:333 in 2xTBS-T were used. Furthermore, instead of *cgl, xylt fucTab* was used for initial depletion of unspecifically binding antibodies contained in the polyclonal HRP mixture.

### Immunoblot analysis of SN *N*-glycoproteins

20μg of protein from SN samples as well as of *A. thaliana* mutant leaf extracts were separated by SDS-PAGE (10 % acrylamide) and transferred to a nitrocellulose membrane. Another aliquot of proteins separated under the same conditions was furthermore stained using Coomassie Blue G. The membrane was blocked with 2 % low-fat milk powder in 1x TBS-T (20 mM Tris, 150 mM NaCl, 0.05 % Tween, pH 7.4) for 16 h at 4 °C and then incubated with affinity purified HRP anti α1,3-fucose and anti β1,2-xylose fractions diluted 1:2 in 2x TBS-T (Kaulfurst-Soboll et al., 2011) for 2 h at RT, respectively. HRP-labeled anti-rabbit antibody (Bio-Rad) was used as secondary antibody 1:10,000 in TBS-T containing 2 % low-fat milk powder for 1 h at RT. Between each step, washing was performed three times with PBS-T. Western blots were developed by enhanced chemiluminescence and signals were digitally recorded with a Fusion-SL imaging system (Peqlab).

### Crossing of IM strains

After mating, mutant strains carrying insertional cassettes in the desired genomic regions were identified by PCR with the respective primers (Table S2). In addition, the mating type of the strains was determined. First, IM*_FucT_* (mt-) was crossed with IM*_Man1A_*xIM*_XylT1A_* (mt+) resulting in IM*_Man1A_*xIM*_XylT1A_*xIM*_FucT_* (mt-). Another strain, created in parallel, was the double mutant IM*_XylT1AB_* (mt+) by mating IM*_XylT1A_* (mt+) and IM*_XylT1B_* (mt-). In order to obtain the quadruple mutant IM*_Man1A_*xIM*_XylT1AB_*xIM*_FucT_*, the triple mutant IM*_Man1A_*xIM*_XylT1A_*xIM*_FucT_* was crossed with the double mutant IM*_XylT1AB_*. In addition to the quadruple mutant, also three other strains: IM*_Man1A_*xIM*_XylT1AB_*, IM*_XylT1A_*xIM*_FucT_* and IM*_XylT1A_*xIM*_FMcT_*xIM*_XylT1B_* were identified in the progenies of the cross. All strains obtained by mating were streaked into single cell colonies, checked again by PCR and then used for mass spectrometry analysis.

### Filter Aided Sample Preparation for *N*-glycoproteomics

Filter Aided Sample Preparation (FASP) was performed as previously described (Mathieu-Rivet et al., 2013, Wisniewski et al., 2009) loading 60 μg protein from SN samples of WT and IM strains onto Amicon ultra centrifugal filters (0.5 mL, 30 kDa MWCO, Millipore) and digesting with 1.2 μg trypsin (sequencing-grade modified, Promega) for 16 h at 37 °C. Peptides were dried in a vacuum centrifuge and stored at −20 °C. Samples from at least three biological replicates of each strain have been analyzed.

### LC-MS analysis

Peptides obtained from FASP were reconstituted in 2 % (v/v) acetonitrile/0.1 % (v/v) formic acid in ultrapure water and separated with an Ultimate 3000 RSLCnano System (Thermo Scientific). The sample was loaded on a trap column (C18 PepMap 100, 300 μm x 5 mm, 5 mm particle size, 100 Å pore size; Thermo Scientific) and desalted for 5 min using 0.05 % (v/v) TFA/2 % (v/v) acetonitrile in ultrapure water with a flow rate of 10 μL min^-1^. Peptides were then separated on a separation column (Acclaim PepMap100 C18, 75 mm i.D., 2 mm particle size, 100 Å pore size; Thermo Scientific) with a length of 50 cm. The mobile phases were composed of 0.1 % (v/v) formic acid in ultrapure water (A) and 80 % acetonitrile/0.08 % formic acid in ultrapure water (B). The gradient used for peptide elution with a flow rate of 300 nL min^-1^ is specified in Table S3.

The LC system was coupled via a nanospray source to a Q Exactive Plus mass spectrometer (Thermo Scientific) operating in positive ion mode. MS data were acquired at a resolution of 70,000 for MS1. Fragmentation by higher-energy C-trap dissociation for MS2 (resolution of 17,500) was triggered in a data-dependent manner dynamically choosing the 12 most abundant precursor ions. Further details for the employed methods are listed in Table S3. However, it should be noted that IS-CID was applied for the analysis of intact *N*-glycopeptides leading to the fragmentation of glycosidic bonds before MS1 (Mathieu-Rivet et al., 2013, Hsiao and Urlaub, 2010). For these measurements, the 12 most abundant precursor ions were chosen by mass tags using masses of +/− 203.079373 (corresponding to the neutral loss of one HexNAc) with charges from 1 - 4 and 5 ppm mass tolerance.

### Identification of peptide spectrum matches (PSMs) and statistical postprocessing

Data analysis was carried out as described in Schulze et al., 2018 employing data base search carried out with MS-GF+ (Kim et al., 2010), X!Tandem (Craig and Beavis, 2003), and OMSSA (Geer et al., 2004) followed by statistical postprocessing of unified results with Percolator (Kall et al., 2007) and qvality (Kall et al., 2009).

### Annotation of *N*-glycan compositions

The annotation of *N*-glycan compositions, employing the in-house developed Python tool SugarPy can be found in Schulze et al., 2017 and Schulze et al., 2018. Therefore, the procedure is summarized here only briefly. The analysis of intact *N*-glycopeptides by IS-CID allows for the identification of the peptide sequence on MS2 level while the *N*-glycan composition can be deduced from a series of neutral losses in the corresponding MS1 spectra. The underlying principle of SugarPy is a creation of all possible combinations for a list of given monosaccharides and a maximal glycan length. In this study, the monosaccharides HexNAc (C8H13NO5), Hex (C6H10O5), MeHex (C7H12O5), dHex (C6H10O4), Pent (C5H8O4) and a maximal glycan length of 15 were used. These combinations are added to potential *N*-glycopeptides identified by use of Ursgal (Kremer et al., 2016) and the resulting library of theoretical glycan tree-peptide combinations are matched on all MS1 spectra employing pyQms (Leufken et al., 2017) for accurate calculation and matching of isotope patterns. For each glycopeptide, a peptide identification (retention time +/− 1 min) was required to accept the identified glycan composition. Finally, when fulfilling certain automatic validations, *N*-glycopeptides are annotated in corresponding MS1 spectra by SugarPy, employing Plotly (Plotly Technologies Inc. Collaborative data science. Montréal, Canada (https://plot.ly) and reviewed manually for the assigned glycan composition. For a better understanding of the matching procedure, exemplary spectra of manually verified *N*-glycopeptides can be found in Figure S3, while all annotated spectra can be found in Supplemental Data 3.

### Calculation of *N*-glycan differences between different strains

Data for bar charts given e.g. in Figures 3, 4 and 5 were created under following considerations. Since using IS-CID following standard peptide database search (on MS2) and *N*-glycan analysis on MS1, *N*-glycan compositions attached to the same peptide can be directly compared between different strains. Therefore, in order to assess certain differences between strains, e.g. the number of dHex attached to *N*-glycopeptides, the number of dHex identified in the *N*-glycan composition attached to peptide X in WT was subtracted from the number of dHex identified in the *N*-glycan composition attached to the very same peptide X in IM*_FucT_*. Considering dHex, the differences can vary from −1 to +1 (as only one dHex is expected being attached to *N*-glycans in *C. reinhardtii)* while for Pent, values between −2 and +2 are occurring. As for many peptides not only one *N*-glycan composition was identified attached to peptide X, but especially *N*-glycan length often deviated between 4 and 8 Hex residues, additional categories were introduced to cover also these microheterogeneities. If e.g. the number of Hex ranged from 6 to 7 in WT (notably still attached to one peptide X) while in IM*_FucT_* only *N*-glycan composition with 5 Hex were identified attached to peptide X, the lowest as well as the highest number of Hex in WT was substracted from IM*_FucT_*. In this example, the *N*-glycosite would be assigned to the category ≤ −1 (as 5-6= −1 and 5-7= −2). In cases of not assignable *N*-glycosites, this range varied across zero (meaning e.g. from −2 to +1).

### Data availability

Mass spectrometry data have been uploaded to the ProteomeXchange Consortium via the PRIDE partner repository (Vizcaino et al., 2013) with the data set identifier PXD012107 for the analysis of intact *N*-glycopeptides experiment. All annotated spectra for the analysis of the *N*-glycan composition can be found in Supplemental Data 3.

## Supporting information

all_supplemntal_figures_and_tables

all_supplemental_information

## Acknowledgments

M.H. acknowledges support from the Deutsche Forschungsgemeinschaft (HI 739/12-1) and by the Sino-German Center, Beijing, China (project GZ990).

## Author contributions

M.H. conceived the idea for the project. A.O, HL and ZZ performed the crossing and, together with M.S., conducted mass spectrometric analysis. A.O and M.H. analyzed the data. S.S. contributed to computational analyses. A.O. wrote the manuscript with S.S. and M.H. M.H. agrees to serve as the author responsible for contact and ensures communication.

---

**Supplemental Data 1**. Peptides used for protein quantification by PRM.

Peptide quantification results conducted by mass spectrometry and evaluated using Skyline for Man1A, XylT1A, XylT1B and the chloroplast ATPase.

**Supplemental Data 2**. Summary of identified *N*-glycan compositions for strains analyzed.

For each *N*-glycopeptide, all identified *N*-glycan compositions are given as separate rows together with the SugarPy output columns, the corresponding proteins (or protein groups with individual proteins separated by ‘<|>‘) as well as the filename and spectrum number of the identification. Strains are separated on different sheets. Annotated spectra for all identified *N*-glycan compositions can be found in Supplemental Data 3.

**Supplemental Data 3**. Annotated spectra for all identified *N*-glycopeptides are sorted by the corresponding strain.

The name of each file includes the name of the corresponding mzML file as well as the *N*-glycopeptide and the spectrum number. Measured peaks are depicted as grey sticks (centroided). Matching isotope patterns of theoretical fragment ions of *N*-glycopeptides are shown as green triangles including a mass accuracy of 10 ppm. Unmatched peaks of the same isotope pattern are shown as red triangles. Annotations are given as blue triangles for the monoisotopic peak. Mouseover on these labels shows all *N*-glycan compositions corresponding to this chemical composition while one of them is also shown above the labeled peak. It should be noted that zooming in and out is supported and recommended.

