## Supplementary material for "Novel insights into *N*-glycan fucosylation and core xylosylation in *C. reinhardtii*": all_supplemntal_figures_and_tables

**Supporting Figures**

Figure S1 Page S-2

Figure S2 Page S-3

Figure S3 Page S-4

Figure S4 Page S-6

Figure S5 Page S-7

Figure S6 Page S-8

Figure S7 Page S-9

Figure S8 Page S-10

Figure S9 Page S-11

**Supporting Tables**

Table S1 Page S-12

Table S2 Page S-13

Table S3 Page S-13

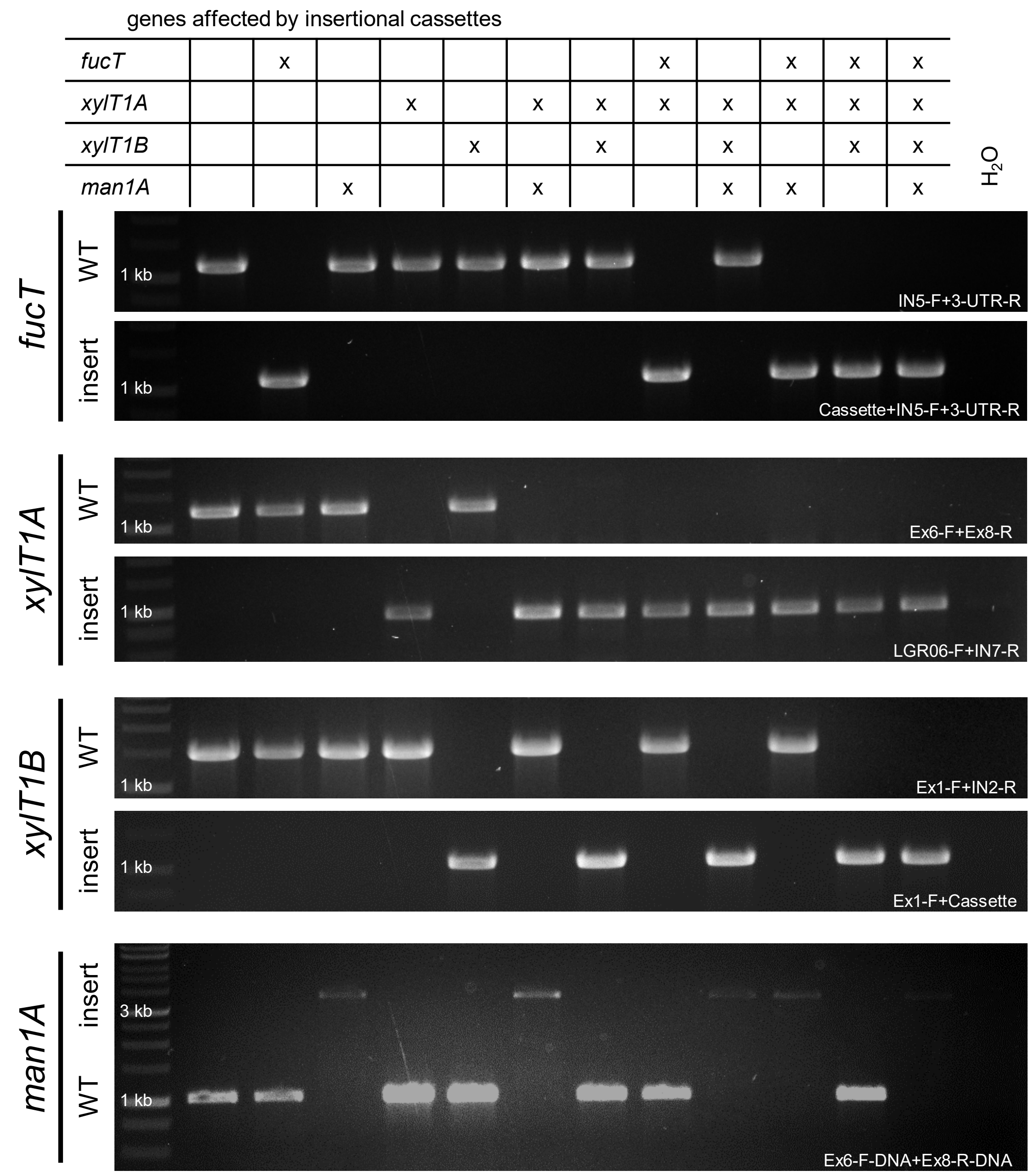


**Figure S1. Verification of *aphVIII* cassettes in respective genes of IM strains analyzed.**

Primer pairs indicated at the right side were used in standard PCR reactions to test, whether genomic WT like regions or insertions of the DNA cassette encoding for the *aphVIII* gene, thus disrupting the coding sequence, are found at the respective sites. 1 % agarose gels stained employing MidoriGreen.

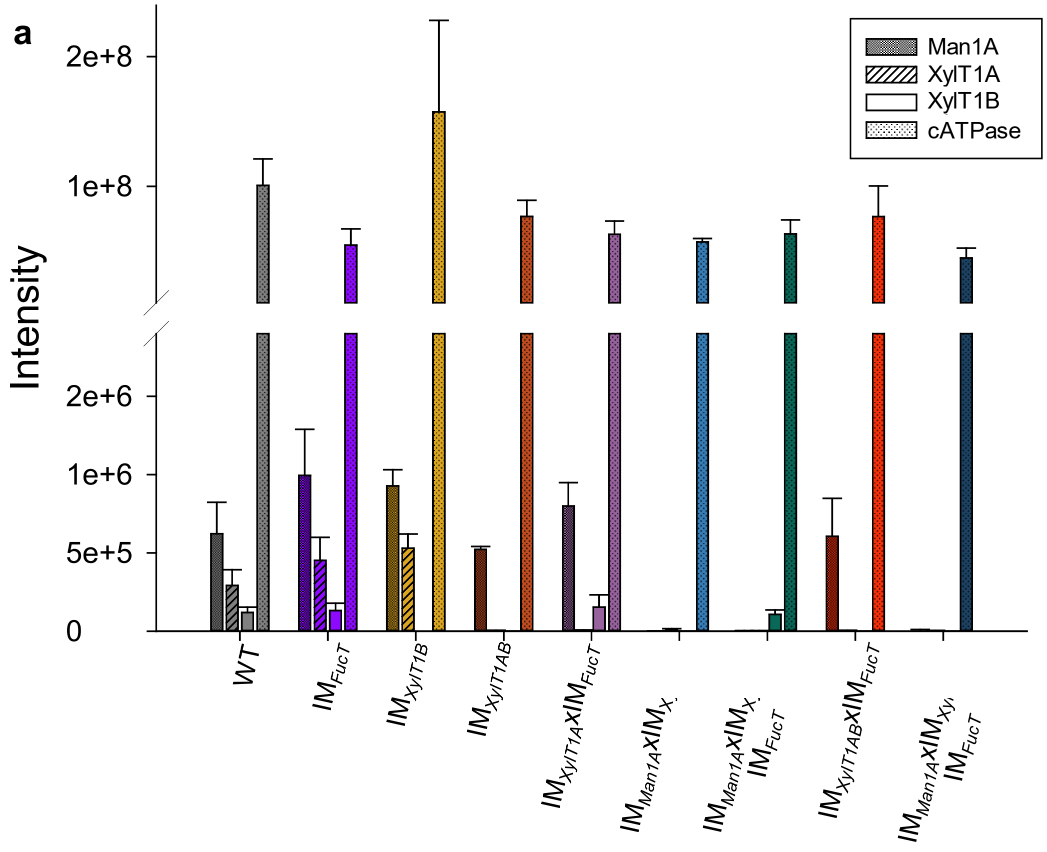


**b**

genes affected by insertional cassettes

| | | | | | | | | | |
|---|---|---|---|---|---|---|---|---|---|
| *fucT* | | x | | x | | x | | x | x |
| *xylT1A* | | | | x | x | x | x | x | x |
| *xylT1B* | | | x | | x | | x | x | x |
| *man1A* | | | | | | x | x | | x |

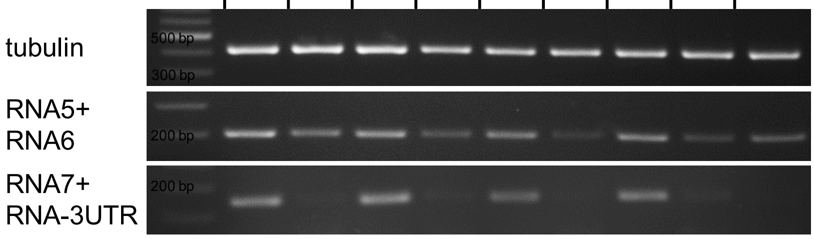


**Figure S2. PRM and cDNA reveal knockdown of affected enzymes in IM strains.**

Parallel Reaction Monitoring (PRM) analyses assessing protein levels of enzymes encoded by genes disrupted due to insertional mutagenesis (Man1A, XylT1A and XylT1B) reveal signals below the detection limit (a). Error bars represent standard deviation between two (including a label swap) biological replicates for PRM results. Since no peptide of FucT was reliably detectable in WT, cDNA analysis was performed for FucT (b). Primer pairs binding prior (RNA5+RNA6) and after the insertion site (RNA7+RNA-3UTR) indicate a knockdown of FucT in the corresponding IM strains. Primer sequences can be found in Table S2.

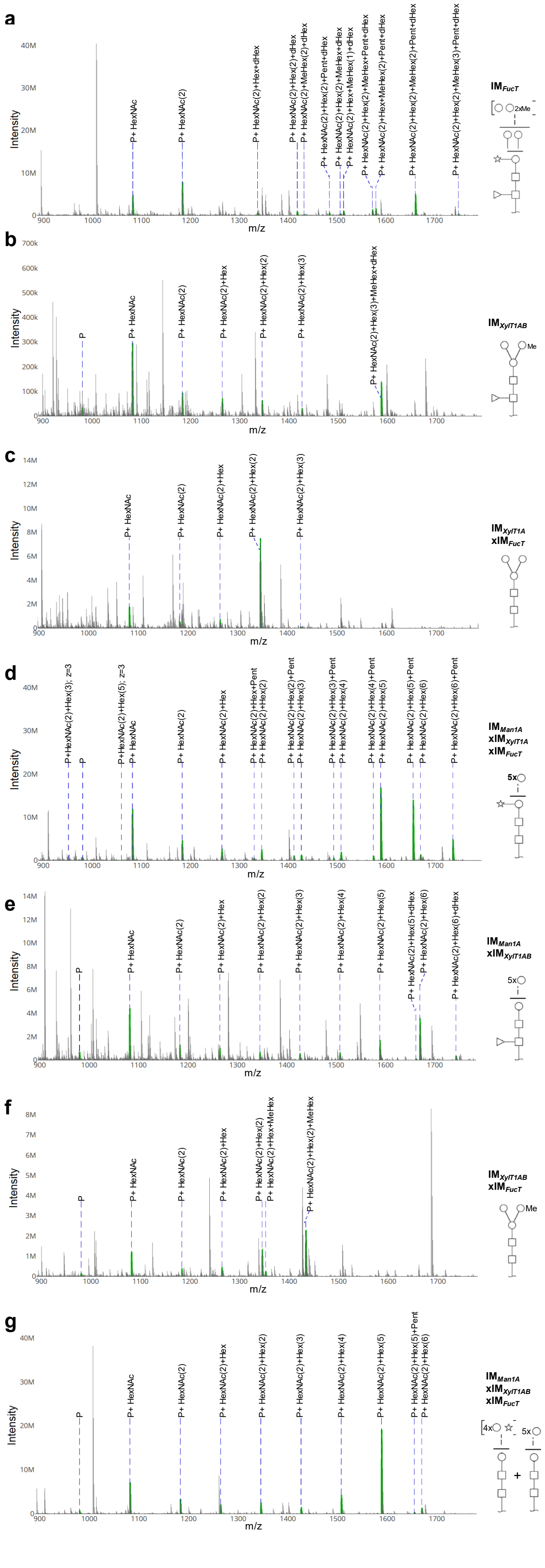

a
IM_FucT
P+ HexNAc
P+ HexNAc(2)
P+ HexNAc(2)+Hex+dHex
P+ HexNAc(2)+Hex(2)+dHex
P+ HexNAc(2)+MeHex(2)+dHex
P+ HexNAc(2)+Hex(2)+Pent+dHex
P+ HexNAc(2)+Hex(2)+MeHex+dHex
P+ HexNAc(2)+Hex+MeHex(1)+dHex
P+ HexNAc(2)+Hex(2)+MeHex+Pent+dHex
P+ HexNAc(2)+Hex+MeHex(2)+Pent+dHex
P+ HexNAc(2)+MeHex(2)+Pent+dHex
P+ HexNAc(2)+Hex(2)+MeHex(3)+Pent+dHex
2xMe
b
IM_XylT1AB
P
P+ HexNAc
P+ HexNAc(2)
P+ HexNAc(2)+Hex
P+ HexNAc(2)+Hex(2)
P+ HexNAc(2)+Hex(3)
P+ HexNAc(2)+Hex(3)+MeHex+dHex
Me
c
IM_XylT1A
xIM_FucT
P+ HexNAc
P+ HexNAc(2)
P+ HexNAc(2)+Hex
P+ HexNAc(2)+Hex(2)
P+ HexNAc(2)+Hex(3)
d
IM_Man1A
xIM_XylT1A
xIM_FucT
P+HexNAc(2)+Hex(3); z=3
P
P+HexNAc(2)+Hex(5); z=3
P+ HexNAc
P+ HexNAc(2)
P+ HexNAc(2)+Hex
P+ HexNAc(2)+Hex+Pent
P+ HexNAc(2)+Hex(2)
P+ HexNAc(2)+Hex(2)+Pent
P+ HexNAc(2)+Hex(3)
P+ HexNAc(2)+Hex(3)+Pent
P+ HexNAc(2)+Hex(4)
P+ HexNAc(2)+Hex(4)+Pent
P+ HexNAc(2)+Hex(5)
P+ HexNAc(2)+Hex(5)+Pent
P+ HexNAc(2)+Hex(6)
P+ HexNAc(2)+Hex(6)+Pent
5x
e
IM_Man1A
xIM_XylT1AB
P
P+ HexNAc
P+ HexNAc(2)
P+ HexNAc(2)+Hex
P+ HexNAc(2)+Hex(2)
P+ HexNAc(2)+Hex(3)
P+ HexNAc(2)+Hex(4)
P+ HexNAc(2)+Hex(5)
P+ HexNAc(2)+Hex(5)+dHex
P+ HexNAc(2)+Hex(6)
P+ HexNAc(2)+Hex(6)+dHex
5x
f
IM_XylT1AB
xIM_FucT
P
P+ HexNAc
P+ HexNAc(2)
P+ HexNAc(2)+Hex
P+ HexNAc(2)+Hex(2)
P+ HexNAc(2)+Hex+MeHex
P+ HexNAc(2)+Hex(2)+MeHex
Me
g
IM_Man1A
xIM_XylT1AB
xIM_FucT
P
P+ HexNAc
P+ HexNAc(2)
P+ HexNAc(2)+Hex
P+ HexNAc(2)+Hex(2)
P+ HexNAc(2)+Hex(3)
P+ HexNAc(2)+Hex(4)
P+ HexNAc(2)+Hex(5)
P+ HexNAc(2)+Hex(5)+Pent
P+ HexNAc(2)+Hex(6)
4x
5x
+
Intensity
m/z

**Figure S3. MS1 spectra assigned for *N*-glycan compositions using SugarPy.**

MS1 spectra for representative *N*-glycan compositions identified in IM strains as being attached to the peptide ITYATTAAAVTNA**N**LSSYK (Asn residue carying HexNAc indicated by bold type letter). Please note that representation of WT was omitted as had been shown already in Schulze et al. 2018. Furthermore, also $IM_{XylT1B}$ is not presented, since its *N*-glycan composition is identical to WT. If not indicated otherwise, *N*-glycopeptides are present in charge state 2. Matched peaks are highlighted in green.

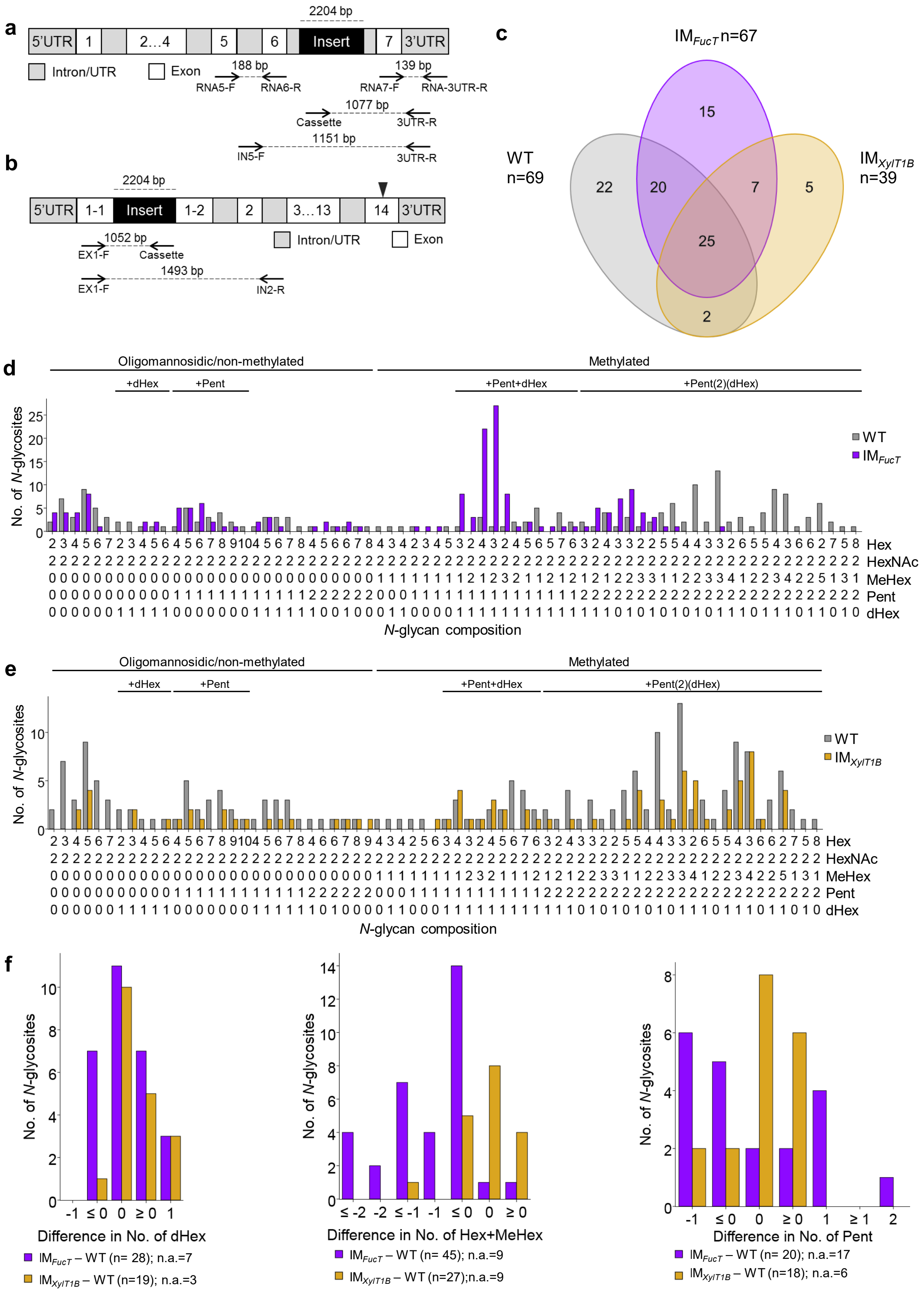


**Figure S4. *N*-glycan analysis of single IM strains reveals only minor changes.**

Schematic representation of the genomic sequence of *fucT* (a) and *xylT1B* (b) (light grey: exon, dark grey: intron, black: insert). Primer pairs surrounding the cassettes were used to verify WT like genomic regions whereas primer pairs involving the cassette primer verified an insertion of the *aphVIII* cassette in the respective genomic regions. The triangle indicate the position of peptides identified by PRM measurements. c, Venn diagram of *N*-glycosites for which the *N*-glycan composition could be determined. Number of all *N*-glycosites harboring the respective *N*-glycan composition are shown for WT (grey), IM$_{FucT}$ (violet) (d) and IM$_{XylT1B}$ (yellow) (e), respectively. The *N*-glycan complexity is increasing from left (oligomannosidic, not methylated) to right (decorated, methylated). *N*-glycan compositions were grouped according to the presence of Pent and/or dHex (optional for sugars written in parenthesis). All *N*-glycosites are taken into account. Peptide sequences and *N*-glycan compositions attached are listed in Supplemental Data 2. f, Differences in the number of dHex (left), *N*-glycan length, defined as the sum of Hex+MeHex (middle), and in the number of Pent (right) for *N*-glycosites found in both strains compared were calculated as depicted in the legend. *N*-glycosites carrying no dHex (left) or Pent (right) in both strains were excluded. The legends indicate the total number of *N*-glycosites compared. Some *N*-glycosites harboring multiple *N*-glycoforms could not be assigned to one of the categories (n.a.).

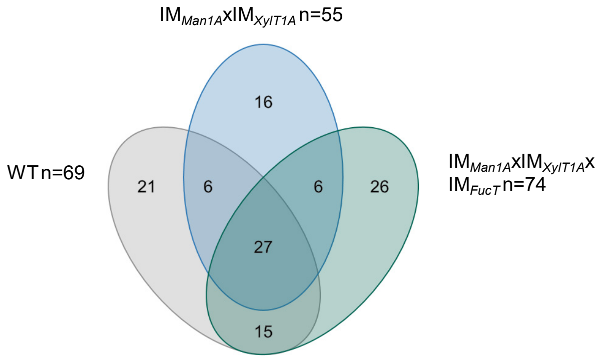


**Figure S5. Analysis of IM$_{Man1A}$xIM$_{XylT1A}$xIM$_{FucT}$**

Venn diagram for *N*-glycosites for which the *N*-glycan composition could be determined in WT, IM$_{Man1A}$xIM$_{XylT1A}$ and IM$_{Man1A}$xIM$_{XylT1A}$xIM$_{FucT}$.

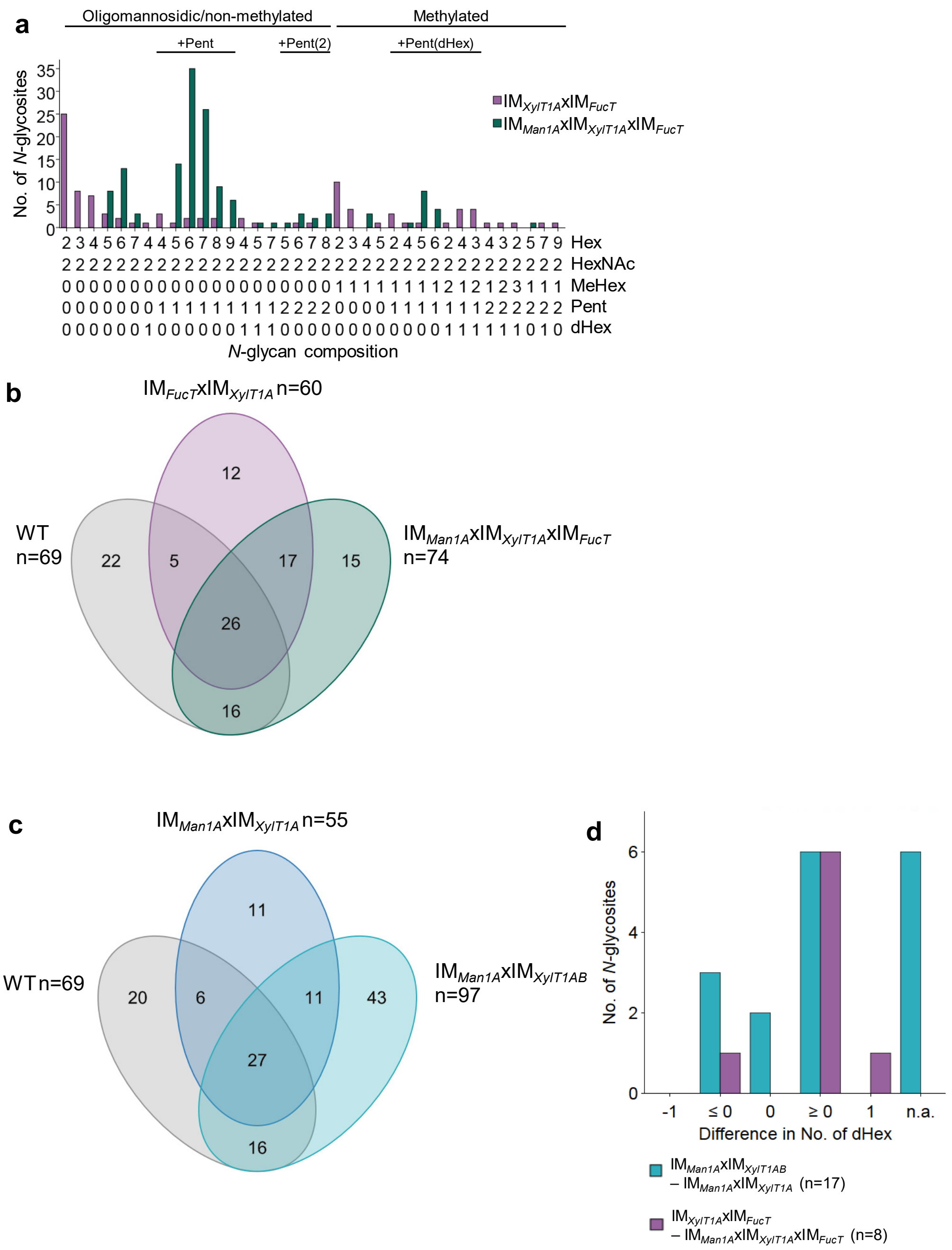


**Figure S6. Analysis of IM*Man1A*xIM*XylT1A*xIM*FucT*, IM*XylT1A*xIM*FucT* and IM*Man1A*xIM*XylT1AB*.**

a, For all identified *N*-glycan compositions, the number of *N*-glycosites harboring this glycan is shown for IM*XylT1A*xIM*FucT* (light violet) and IM*Man1A*xIM*XylT1A*xIM*FucT* (green). The *N*-glycan complexity is increasing from left (oligomannosidic, not methylated) to right (decorated, methylated). *N*-glycan compositions were grouped according to the presence of Pent and/or dHex (optional for sugars written in parenthesis). All *N*-glycosites are taken into account. Peptide sequences and *N*-glycan compositions attached are listed in Supplemental Data 2. b, Venn diagram for *N*-glycosites for which the *N*-glycan composition could be determined for IM*XylT1A*xIM*FucT* and IM*Man1A*xIM*XylT1A*xIM*FucT*. C, Venn diagram for *N*-glycosites for which the *N*-glycan composition could be determined for IM*Man1A*xIM*XylT1A* and IM*Man1A*xIM*XylT1AB*. d, Differences in number of dHex for *N*-glycosites found in both strains were calculated. The legends indicate the total number of *N*-glycosites compared. Some *N*-glycosites harboring multiple *N*-glycoforms could not be assigned to one of the categories (n.a.).

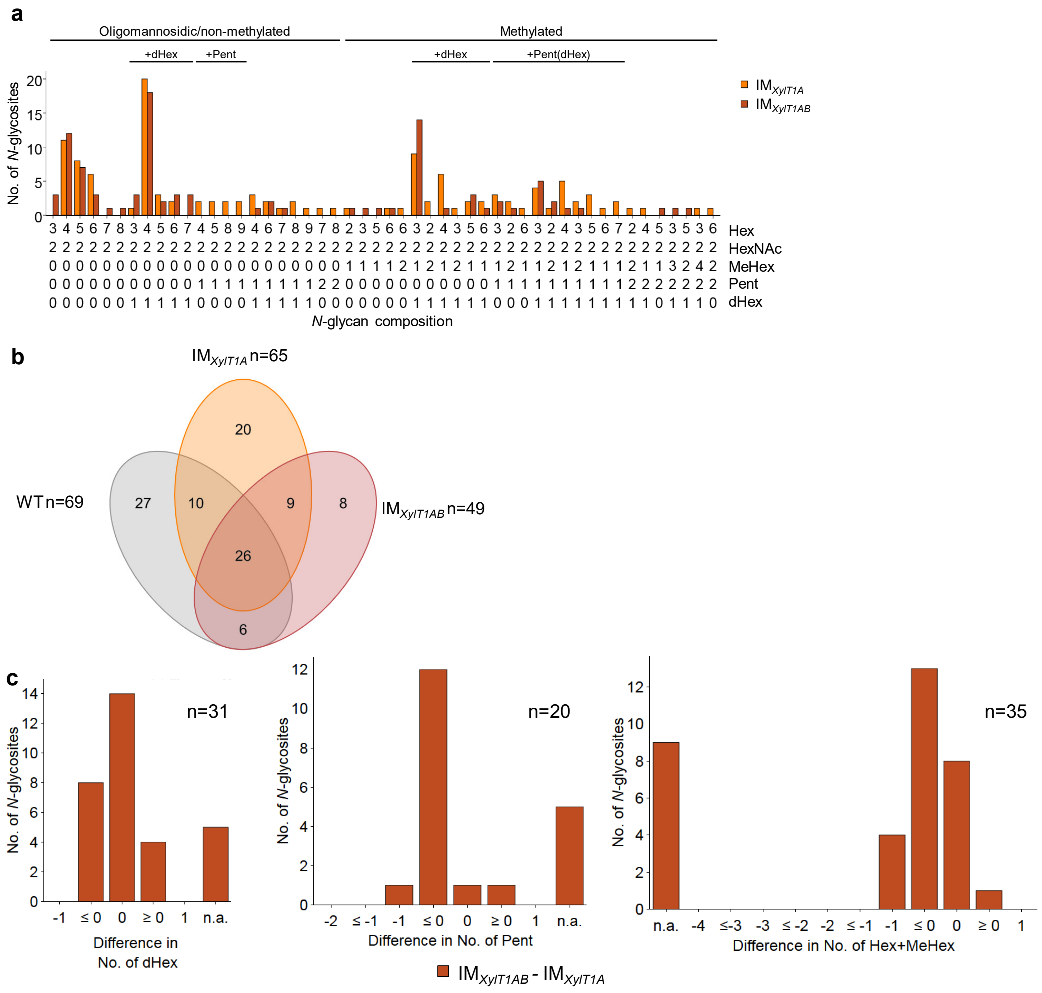


**Figure S7. Analysis of IM$_{XylT1AB}$.**

For all identified *N*-glycan compositions, the number of *N*-glycosites harboring this glycan is shown for IM$_{XylT1A}$ (light orange) and IM$_{XylT1AB}$ (brown) (a). The *N*-glycan complexity is increasing from left (oligomannosidic, not methylated) to right (decorated, methylated). *N*-glycan compositions were grouped according to the presence of Pent and/or dHex (optional for sugars written in parenthesis). All *N*-glycosites are taken into account. Peptide sequences and *N*-glycan compositions attached are listed in Supplemental Data 2. b, Venn diagram for *N*-glycosites for which the *N*-glycan composition could be determined. c, Differences in the number of dHex (left), Pent (middle) and Hex+MeHex (right) for *N*-glycosites found in both strains. *N*-glycosites, carrying no dHex (left) or Pent (middle) in both strains were excluded. The legends indicate the total number of *N*-glycosites compared. Some *N*-glycosites harboring multiple *N*-glycoforms could not be assigned to one of the categories (n.a.).

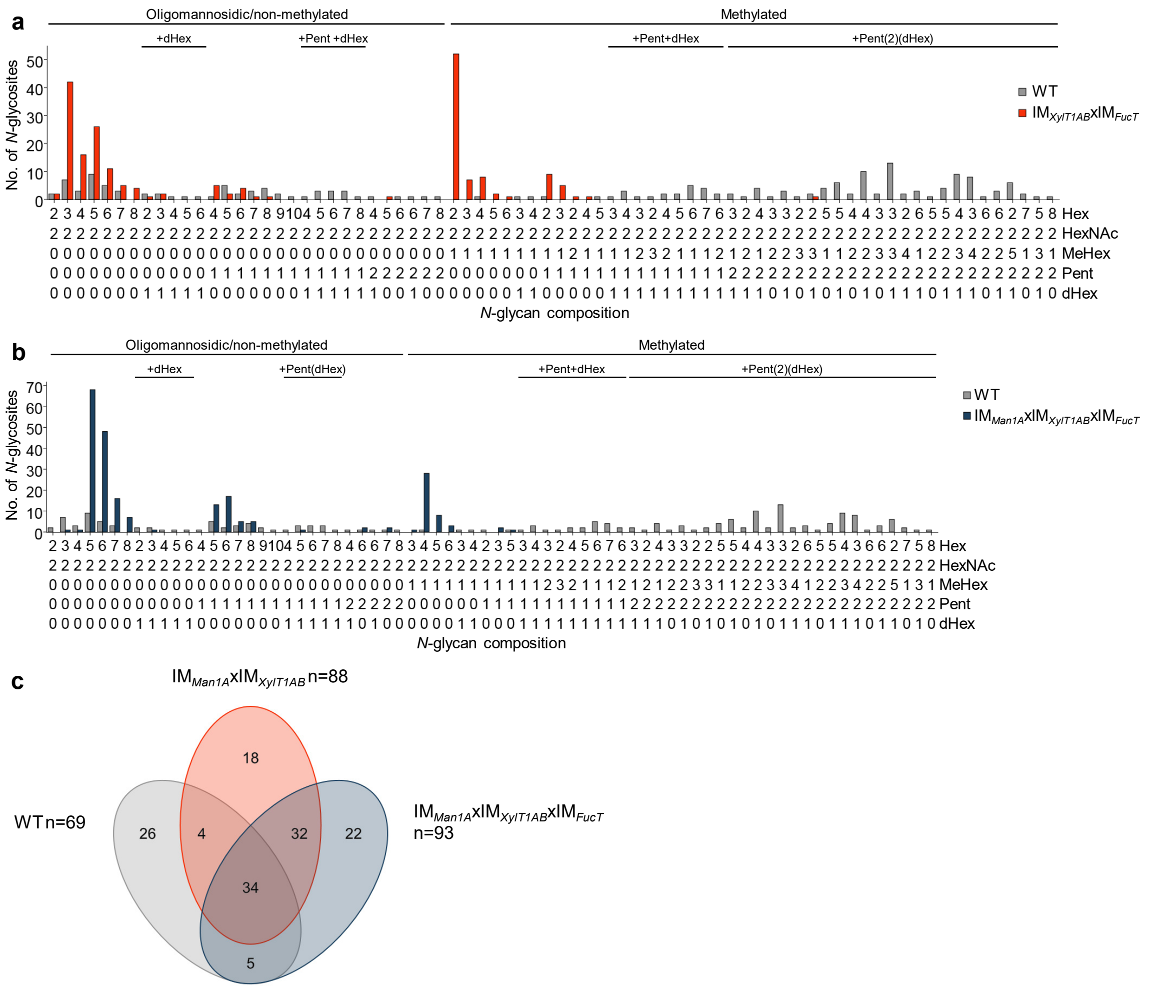


**Figure S8. Analysis of all *N*-glycosites in IM$_{XylT1AB}$xIM$_{FucT}$ and IM$_{Man1A}$xIM$_{XylT1AB}$xIM$_{FucT}$ reveals lack of *N*-glycan core modifications.**

For all identified *N*-glycan compositions, the number of *N*-glycosites harboring this glycan is shown for WT (grey), IM$_{XylT1AB}$xIM$_{FucT}$ (orange) (a) and IM$_{Man1A}$xIM$_{XylT1AB}$xIM$_{FucT}$ (dark blue) (b), respectively. The *N*-glycan complexity is increasing from left (oligomannosidic, not methylated) to right (decorated, methylated). *N*-glycan compositions were grouped according to the presence of Pent and/or dHex (optional for sugars written in parenthesis). All *N*-glycosites are taken into account. Peptide sequences and *N*-glycan compositions attached are listed in Supplemental Data 2. c, Venn diagram for *N*-glycosites for which the *N*-glycan composition could be determined.

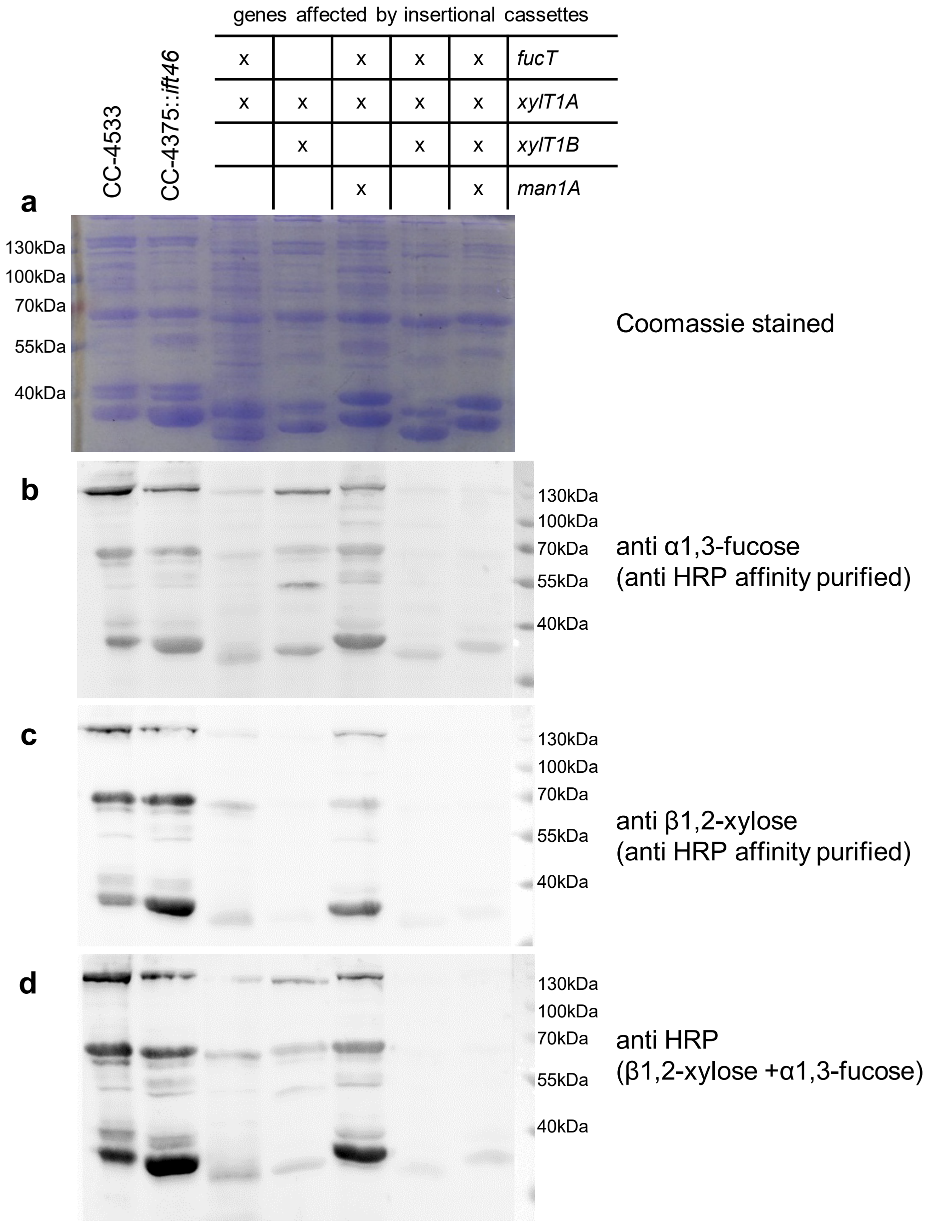


**Figure S9. Immunoblotting of residual IM confirms interdependence of core-xylosylation and fucosylation.**

20 µg of SN proteins of selected IM strains were separated by SDS-PAGE and transferred to nitrocellulose membranes. Additionally, one SDS PAGE gel was stained as loading control using Coomassie Brilliant Blue G (a). Membranes were incubated in affinity purified HRP antibody binding to α-1,3-fucose (b) and β1,2-core xylose (c), respectively, as well as with the polyclonal HRP antibody (d).

**Table S1.** *N*-glycan compositions found in different strains analyzed for methylation degree as well as for Pent and dHex amount.

All *N*-glycosites identified in the strains listed were compared regarding methylation, Pent and dHex numbers attached. Boxes in dark grey highlight the loss of Pent upon excessive *N*-glycan trimming, due to a lack of XylT1-B action. *core Pent is defined as Pent, where the fragment peak Hex(1)HexNAc(2)Pent(1) was matched on MS1 level.

| | **WT** | **$IM_{FucT}$** | **$IM_{XylT1\text{-}A}$x $IM_{FucT}$** | **$IM_{Man1A}$x $IM_{XylT1\text{-}A}$x $IM_{FucT}$** | **$IM_{Man1A}$x $IM_{XylT1\text{-}AB}$** | **$IM_{XylT1\text{-}AB}$x $IM_{FucT}$** | **$IM_{Man1A}$x $IM_{XylT1\text{-}AB}$x $IM_{FucT}$** |
|---|---|---|---|---|---|---|---|
| total No. of *N*-glycosites | 194 | 184 | 122 | 146 | 249 | 225 | 238 |
| *N*-glycosites with >1 MeHex | 124 | 125 | 49 | 18 | 61 | 93 | 48 |
| *N*-glycosites with 1 or 2 Pent | 155 | 157 | 47 | 119 | 36 | 39 | 48 |
| *N*-glycosites with core Pent * | >97 | >97 | >23 | >65 | >2 | >9 | >10 |
| *N*-glycosites with dHex | 123 | 116 | 18 | 3 | 63 | 5 | 2 |
| *N*-glycosites with >1 MeHex / total No. of *N*-glycosites | 63.9% | 67.9% | 40.2% | 12.3% | 24.5% | 41.3% | 20.2% |
| *N*-glycosites with 1 or 2 Pent / total No. of *N*-glycosites | 79.9% | 85.3% | 38.5% | 81.5% | 14.5% | 17.3% | 20.2% |
| *N*-glycosites with 1 core Pent / total No. of *N*-glycosites | >50.0% | >52.7% | >18.9% | >44.5% | >8.0% | >4.0% | >4.2% |
| *N*-glycosites with dHex / total No. of *N*-glycosites | 63.4% | 63.0% | 14.8% | 2.1% | 25.3% | 2.2% | 0.84% |

**Table S2.** Primer sequences used for verification PCRs and mRNA analysis.

Primer sequences used to analyze the presence or absence of an insert in the specific genomic region. Primers used for the verification of an insertion in the genomic regions encoding for Man1A and XylT1-A can be found in Schulze et al. 2018.

In addition, primer sequences used for mRNA analysis of FucT expression levels are given.

| Primer name | Primer sequence | purpose |
|---|---|---|
| EX1-F | CTGTCGGTCGCGCAAATTGG | verification of insert in XylT1-B on DNA level |
| IN2-R | CCACCTCCACATTACGGCCT | |
| IN5-F | TGGAAGCCAGGACAACGCAC | verification of insert in FucT on DNA level |
| 3UTR-R | CACGTCACAGCTTCCTGCCT | |
| Cassette | GCCCACGGTCAATTAGCCAC | verification of insert in XylT1-B and FucT on DNA level |
| RNA5-F | CAAGATGGAGCTGATCCGCG | relative quantification of mRNA level of FucT before insert |
| RNA6-R | AGCCCAGCCTGTTGTAGTCG | |
| RNA7-F | CTTCAGCACCTGCCTGTTCG | relative quantification of mRNA level of FucT following insert |
| RNA-3UTR-R | CTTCAGCACCTGCCTGTTCG | |
| tubulin-F | CACATCCAGGGTGGCCAG | relative quantification of tubulin mRNA (as housekeeping gene) |
| tubulin-R | CCTGGAAGCCCTGCAGGC | |

**Table S3**. List of mass spectrometry related parameters that used for the analysis intact *N*-glycopeptides using IS-CID and PRM measurements.

For HPLC, the mobile phases were composed of 0.1 % (v/v) formic acid in ultrapure water (A) and 80 % acetonitrile/0.08 % formic acid in ultrapure water (B). Percentage of buffer B is indicated as well as its increase (↑) or decrease (↓).

| Method / Parameter | IS-CID for intact *N*-glycopeptides 50 cm column | PRM measurement for protein quantification |
|---|---|---|
| HPLC gradient | 5 min 2.5 %<br>40 min ↑ to 45 %<br>5 min ↑ to 99 %<br>20 min 99 %<br><br>5 min ↓ to 2.5 %<br>30 min 2.5 % | 5 min 2.5 %<br>65 min ↑ to 18 %<br>50 min ↑ to 32 %<br>5 min ↑ to 99 %<br>20 min 99<br>5 min ↓ to 2.5 %<br>30 min 2.5 % |
| IS-CID | 80 eV | - |
| Mass tags | 5 ppm | - |
| Excluded charges | unassigned, ≥5 | - |
| Dynamic exclusion | 15 s | - |
| Lock masses | off | best |
| MS1 scan range | 600 – 3000 *m/z* | 350-1600 *m/z* |
| MS1 AGC target | $3e^6$ | $3e^6$ |
| MS1 maximum injection time | 100 ms | 50 ms |
| MS2 scan range | dynamic, fixed first mass150 *m/z* | dynamic, fixed first mass 100 *m/z* |
| MS2 AGC target | $1e^5$ | $5e^4$ |
| MS2 maximum injection time | 120 ms | 240 ms |
| NCE | 30 | 27 |
